# Dietary inclusion of nitrite-containing frankfurter exacerbates colorectal cancer pathology, alters metabolism and causes gut dybiosis in APC^min^ mice

**DOI:** 10.1101/2022.03.17.484445

**Authors:** William Crowe, Xiaobei Pan, James Mackle, Adam Harris, Gary Hardiman, Chris T Elliott, Brian D Green

## Abstract

Colorectal cancer (CRC) is the second most prevelant malignancy in Europe and diet is an important modifiable risk factor. Processed meat consumption, including meats with preservative salts such as sodium nitrite, have been implicated in CRC pathogenesis. This study investigated how the CRC pathology and metabolic status of adenomatous polyposis coli (APC) multiple intestinal neoplasia *(min*) mice was perturbed following 8 weeks of pork meat consumption. Dietary inclusions (15%) of either nitrite-free pork, nitrite-free sausage or nitrite-containing sausage (frankfurter) were compared against a parallel control group (100% chow). Comprehensive studies investigated: gastrointestinal tract histology (tumours, aberrant crypt foci (ACF) and mucin deplin foci (MDF), lipid peroxidation (urine and serum), faecal microbiota and serum metabolomics (599 metabolites). After 8 weeks mice consuming the frankfurter diet had 53% more (P=0.014) gastrointestinal tumours than control, although ACF and MDF did not differ. Urine and serum lipid peroxidation markers were 59% (P=0.001) and 108% (P=0.001) higher, respectively in the frankfurter group. Gut dysbiosis was evident in these mice with comparably fewer *Bacteriodes* and more *Firmicutes*. Fasting serum levels of trimethylamine N-oxide (TMAO) and numerous triglycerides were elevated. Various serum phosphotidylcholine species were decreased. These results demonstrate that nitrite-containing sausages may exaccerbate the development of CRC pathology in APC^Min^ mice to a greater extent than nitrite-free sausages, and this is associated with greater lipid peroxidation, wide-ranging metabolic alternation and gut dysbiosis.

## Introduction

Colorectal cancer (CRC) is the third most prevalent malignancy worldwide. Over 1.9 million people were diagnosed with CRC in 2020 ^1^. CRC always begins in the form of abnormal growths called polyps in the colon or rectum. The majority of colorectal tumours are adenocarcinomas that originate from polyps. Polyps are pedunculated masses of tissue that may be pre-cancerous. Around 5% of CRC is attributable to familial adenomatous polyposis^2^. The remaining cases of CRC are believed to be influenced by environmental factors. One modifiable risk factor that has been proposed as a leading candidate in the search for causes of CRC is dietary pattern. A diet high in processed meat is suggested as a major risk factor for CRC development. The International agency for Reseach on Cancer (IARC) have advised consumers to avoid consuming processed meat and limit the quantity of red meat consumed.

Processed meat is classed as a group 1 carcinogen, and therefore carcinogenic to humans. This conclusion is based on 9 case-control studies and 18 cohort studies and of these 6 and 12, respectively, linked processed meat consumption to CRC 3^3^.

However, these studies only represent 10% of the existing body of evidence, the remainder of which was deemed uninformative due to methodological limitations. IARC list examples of processed meat as hot dogs (frankfurters), ham, sausages, corned beef, and biltong or beef jerky as well as canned meat and meat-based preparations and sauces. Processed meat is often preserved with nitrite salts, and recent scientific evidence suggests that it is this component that is responsible for processed meats unfavourable link with CRC ^4^. Sodium nitrite uniquely prevents the growth of *Clostridium Botulinum*, it also contributes to the colour and taste of processed meat. Ingested nitrite has the potential to form N-Nitroso compounds (NOC) including nitrosamines, some of which are known to be carcinogenic. NOCs have been proposed to be involved in the aetiology of several types of cancer, particularly those in the gastrointestinal tract ^5^.

The composition of gut microbiota (consisting of more than 10^13^ bacteria), is modulated by exposure to environmental factors such as diet and medication. These microbes protect the host by inhibiting the growth of pathogens and contributing to the production of short chain fatty acids (SCFA). SCFAs include acetate, butyrate, and propionate are used an energy source and participate in the proliferation of intestinal epithelial cells^6^. Dysbiosis has been implicated in the pathogenesis of numerous health conditions, including CRC ^7^. It has been reported in CRC that there is a depletion of *Bacteroidetes* and *Firmicutes*, and at the order level there is a reduction in *Clostridiales* ^8^. Sodium nitrite is also known to prevent the colonisation of *Clostridiales*.

A total of ten animal studies have investigated the effect of nitrite consumption on CRC, 4 of these studies found that nitrite consumption had no effect on CRC, 5 studies found that nitrite consumption significantly increased the risk of CRC, and 1 study found that nitrite containing processed meat significantly decreased the risk of CRC^4^. All studies reporting a significant causative effect supplied ≥50% processed meat in the diet. This does not reflect typical human consumption and therefore further studies employing more realistic levels of processed meat are required.

Current public health advice treats all processed meat equally, when in reality there are many differences in their composition. Most notably there is the addition of sodium nitrite as a preservative which is present in some products and not in others. For example, nitrites are not traditionally an ingredient in the manufacture of traditional British/Irish sausages but it is more common in continental European sausages. Therefore, the primary aim of this investigation was to examine whether cancer progression in a mouse model of CRC pathology differs when fed a moderate amount of either nitrite-containing frankfurter sausage or nitrite-free sausage. A secondary aim was to examine whether CRC development was impacted by the consumption of either processed (nitrite-free sausage) or unprocessed (pork) meat. A number of additional analyses were undertaken to ascertain what the wider metabolic consequences of these meat diets.

## Results

### Intestinal pathology

In the duodenum the mean tumour count for mice fed a 15% frankfurter diet was 4.2 (Figure 1b). This was 75% higher than the mean tumour count of the control group (p=0.022). No other inter-group differences were detected for duodenal tumours. The quantity of tumours in the jejunum did not differ between any groups. Frankfurter fed animals had 88.2% more tumours in their ilieum than the control (p=0.011), and 190.9% more than pork fed mice (p=0.001), but did not significantly different from the nitrite-free sausage group. Sausage fed mice had 154.6% more tumours in their ilieum than the pork fed mice (p=0.050). In the colon, the only significant difference was that pork fed mice had 81.8% more tumours than the control mice (p=0.047). In terms of the overall number of GI tumours, the frankfurter group had the highest with a mean of 11.2 and this was significantly more (p=0.002) than the control which had a mean of 7.3 tumours, it was also significantly higher (p=0.029) than the sausage group which had a mean of 8.6 tumours, and significantly higher (p=0.019) than the pork group which had a mean of 8.3 tumours.

**Figure 1:**
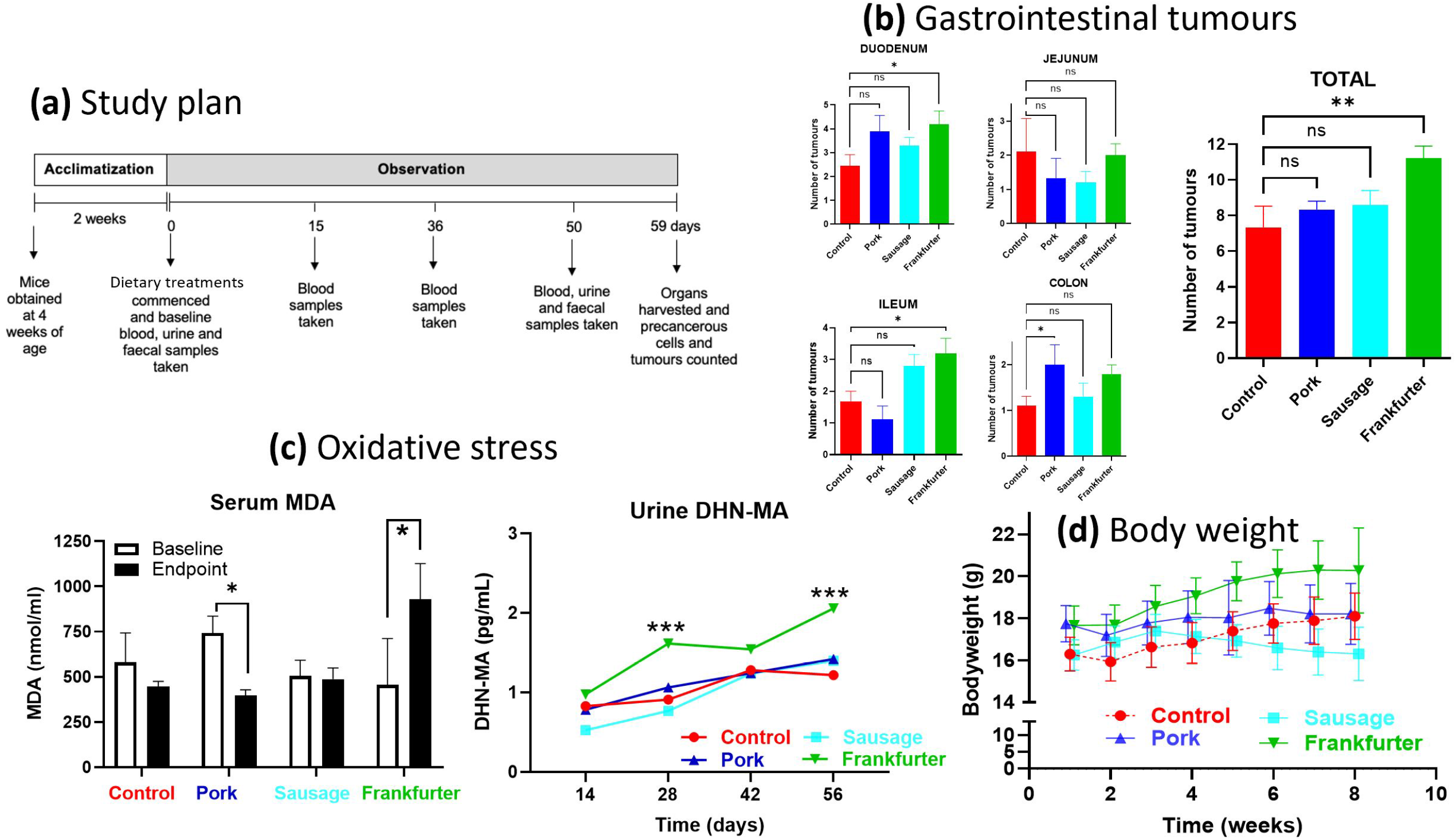
APC^Min^ mice consuming 15% frankfurter (85% chow) for 8 weeks have significantly more gastrointestinal tumours, higher levels of lipid peroxidation, and greater body weight than APC^Min^ mice consuming a control diet (100% chow). **a)** after acclimatisation mice were fed one of four chow diets which were either Control (n=9; standard chow (red)), or chow with a 15% incorporation of pork mince (n=9; blue), pork sausage (n=10; light blue), or nitrite-containing frankfurter sausage (n=10; green). **b)** numbers of tumours did not differ between any groups in any of the specific regions of the GI tract. However, compared with control signifcantly more tumours were displayed by the frankfurter group alone. **c)** terminal serum MDA concentrations of mice consuming frankfurter were elevated compared to that of control terminal samples, after controlling for baseline using an ANCOVA. Urine DHN-MA concentrations were compared between groups at the same timepoint using an ANOVA. Levels in mice consuming frankfurter supplemented chow had higher levels at 28 and 56 days. **d)** bodyweights of frankfurter-fed mice were consistently higher than control mice. Error bars shown represent s.e.m.

As shown in Table 1 no significant differences were found in ACF or MDF between any dietary group. There was a trend for pork-fed mice to have the highest quantities of ACF (not significant), and for sausage-fed mice to have the highest quantities of MDF (not significant).

**Table 1.** Numbers of aberrant crypt foci and mucin depleted foci did not significantly differ between any groups.

|  | **Control** | **Pork** | **Sausage** | **Frankfurter** |
| --- | --- | --- | --- | --- |
| **ACF** | 102.44 (10.86) | 115.11 (17.03) | 101.80 (10.71) | 101.90 (15.18) |
| **MDF** | 6.11 (2.76) | 5.98 (2.67) | 6.5 (2.59) | 5.08 (3.12) |

ACF- aberrant crypt foci; MDF- mucin depleted foci. Mean values were compared between groups by analysis of variance, alpha value of less than 0.05 was accepted as statistically significant. Values in parenthesis are standard deviations for the corresponding mean.

### Lipid peroxidation

Frankfurter feeding consistently led to the highest concentrations of urinary DHN-MA (Figure 1c). By week 3 these were 54.30% higher than control (p=0.001), 47.39% higher than sausage-fed (p=<0.001), and 65.66% higher than pork-fed (p=<0.001). By week 3 Pork-fed mice had 72.17% higher urinary DHN-MA than sausage-fed (p=0.048). By week 6 the frankfurter-fed group had DHN-MA levels that were 83.05% higher than the control (p=0.030), 80.26% higher than sausage-fed (p=0.009), and 80.66% higher than pork-fed (p=0.013). At week 8 the frankfurter group had DHN-MA levels that were 59.18% higher than the control (p=0.001), 67.92% higher than sausage-fed (p=<0.001), and 69.07% higher than pork-fed (p=0.001).

Similar results were found when comparing serum MDA concentrations between groups. Frankfurter-fed mice had the highest mean concentration (928.89 pg/ml) and this was 48.07% higher than the control (p=<0.001), 42.90% higher than pork-fed (p=<0.001), and 52.56% higher than sausage-fed (p=0.001). These were all significantly different after controlling for mouse baseline values.

### Bodyweight and food intake

As shown in Figure 1d, mice fed pork or frankfurter diet had significantly higher body weight than the control and sausage groups at week 1. At almost every timepoint the frankfurter group had the highest mean bodyweight and this was significantly higher than the control. At weeks 6, 7, and 8 the sausage group had significantly lower mean bodyweights than the control. At weeks 2-6 the pork group had significantly higher mean bodyweights than the control, but not at weeks 7 and 8.

### Altered concentrations of serum metabolites

Unsupervised Principal Component Analysis (PCA) of serum metabolomic data explained 89.4% of the total variation (76.9%, 8.0% and 4.5% for PC1, PC2 and PC3, respectively; Figure 2a). There was visible separation with a tendency for frankfurter group profiles to separate from the other three groups along the PC1 axis. Profiles from control, pork and sausages groups were more closely clustered together with variation more restricted to the PC2 axis.

**Figure 2.**
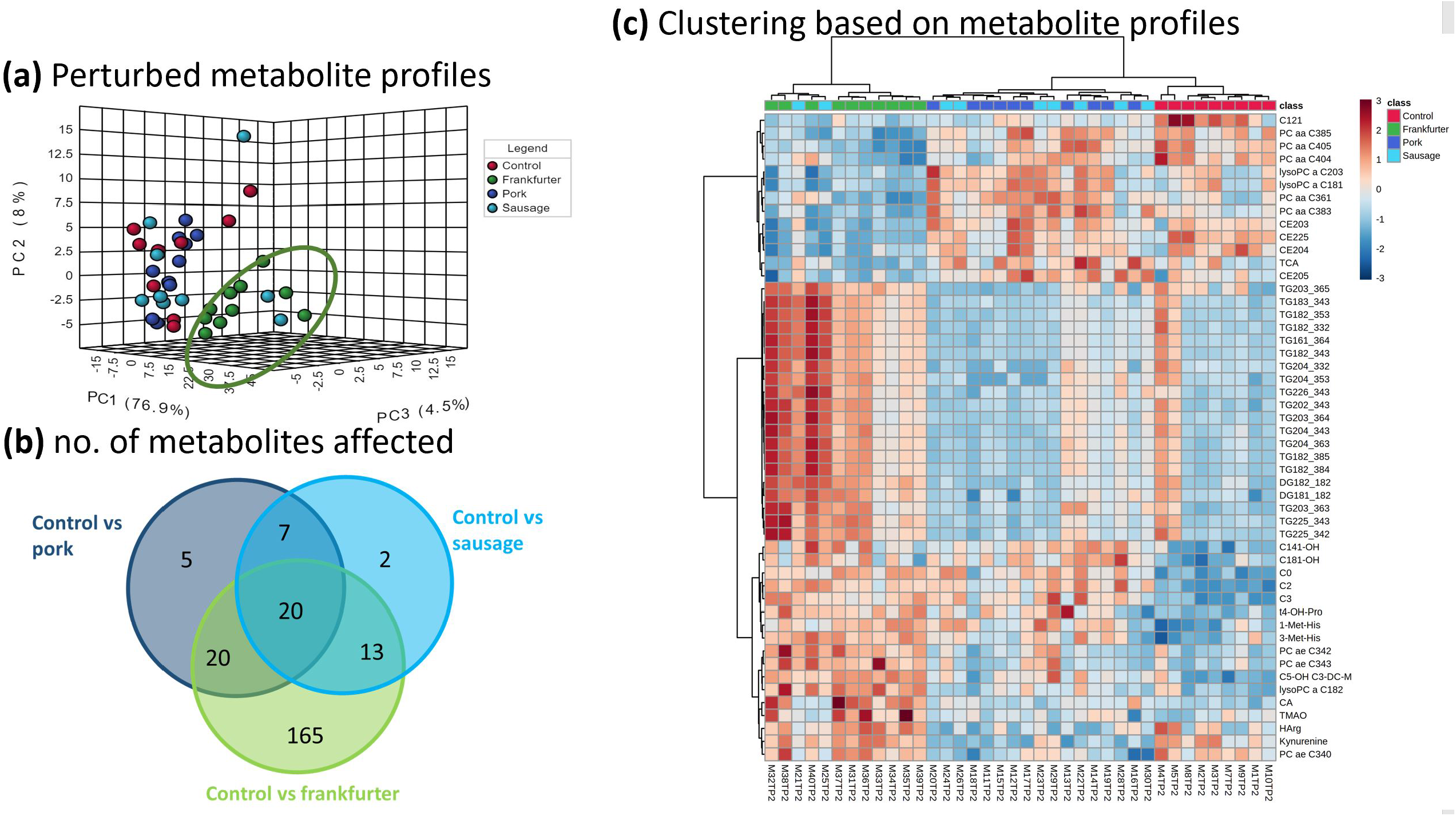
Serum metabolite profiles of frankfurter-fed mice deviate most from control, whereas those fed pork or sausage align more closely. Serum collected at 56 days under went metabolomic profiling: **a)** shows scores plots of unsupervised PCA based on the first three principle components (PCs) for Control (n=9; standard chow (red)), or chow with a 15% incorporation of pork mince (n=9; blue), pork sausage (n=10; light blue), or nitrite-containing frankfurter sausage (n=10; green). Metabolomic data explained 89.4% of the total variation (76.9%, 8.0% and 4.5% for PC1, PC2 and PC3, respectively). There was a tendency for profiles from the frankfurter group to separate along the PC1 axis. Profiles from control, pork and sausages groups clustered more closely with variation more restricted to PC2. **b)** is a Venn diagram showing the number of the 599 metabolites measured which significantly differed in group comparisons with control. c) heat map with hierarchical clustering based on the top 50 metabolites ranked by FDR –corrected P-value (ANOVA). Each row represents a metabolite and each column a sample. Red (high) and blue (low) indicates relative concentrations. Dendrograms indicating similarity are shown at top (samples) on left side (metabolites). Serum metabolite measurements readily clustered mice by diet with pork and nitrite-free sausage being most similar to one another. Frankfurter consumption was characterised by significantly higher levels of certain species of diglycerides (DG), triglycerides (TG) and acylcarnitines (C), and by lower levels of certain phosphatidylcholines (PC/LysoPC) and cholesterol esters (CE). Other notable observations included the differing bile acid responses of taurocholic acid (TCA) and cholic acid (CA). The profound variability of trimethylamine N-oxide (TMAO) responses in frankfurters was also very evident. Control samples are shown in red, minced pork in dark blue, nitrite-free sausage in light blue, and frankfurter (nitrite-containing) in green.

Serum metabolites were ranked by false-discovery rate (FDR) corrected p-values based on one-way ANOVA. Out of 599 measured metabolites 232 significantly differed between two or more groups. When each of the dietary interventions was compared with Control it was the frankfurter diet which caused by far the greatest number of metabolite alternations (Figure 2b). In total 218 metabolite alternations were observed for frankfurter (165 of which were unique), whereas comparably fewer were observed for pork and sausage, 52 and 42, respectively. Hierarchical clustering analysis (HCA) was conducted by selecting the 50 most statistically significant metabolites from the ANOVA found that the greatest distinction in profiles was between the control and frankfurter-fed mice (Figure 2c). Metabolic profiles obtained for pork-fed and sausage-fed mice were similar to one another and were clustered together by HCA. From this it was evident that many triglyceride species were significantly higher in the frankfurter group, and several certain cholestrol esters and phosphotidylcholines were significantly lower.

Figure 3 shows that serum TMAO (trimethylamine N-oxide) levels were 184% higher in mice receiving frankfurter diet. However, the levels of TMAO precursor metabolites (choline, betaine, carnitine, acetylcarnitine propionylcarnitine) are not similarly affected. Levels of choline and betaine did not differ between groups. Mice consuming any of the meat-containing diets had similarly higher levels of carnitine (71-119%), acetylcarnitine (114-217%) or propionylcarnitine (97-155%). In no cases were these carnitine metabolites the highest overall in mice consuming the frankfurter diet.

**Figure 3.**
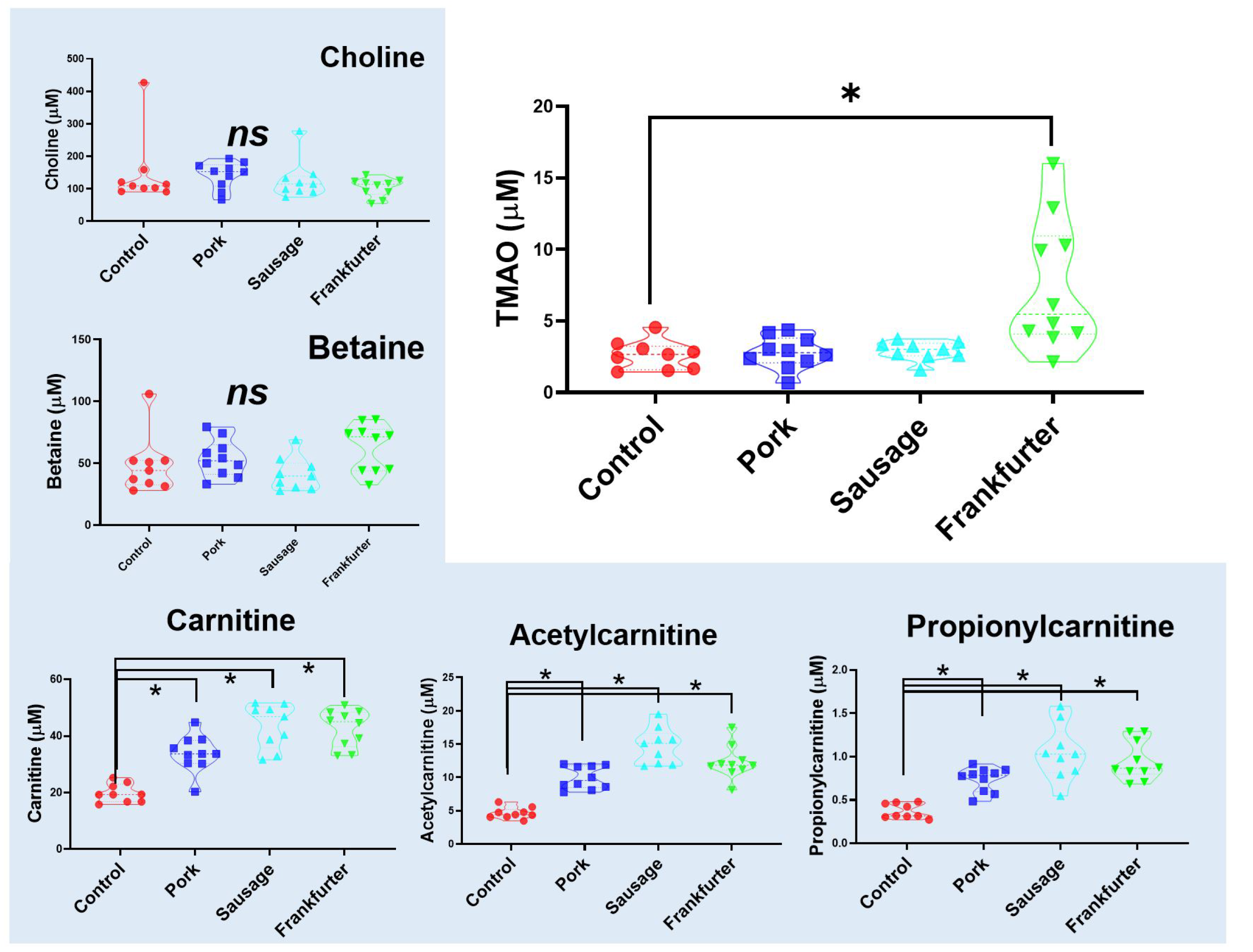
Serum TMAO (trimethylamine N-oxide) is higher and more variable in mice receiving frankfurter diet, yet TMAO precursor metabolites are not similarly affected. Serum levels of TMAO (at 56 days), typically formed from choline, betaine and carnitines were significantly higher in mice receiving frankfurter-containing chow. Levels of choline and betaine did not differ between groups. Mice consuming any meat-containing chow had higher levels of carnitine, acetylcarnitine, and propionylcarnitine. Control samples are shown in red (n=9), minced pork in dark blue (n=10), nitrite-free sausage in light blue (n=9), and frankfurter (nitrite-containing) in green (n=10). Data shown are violin plots containing individual samples values with the median and quartile values represented as dotted lines.

### Faecal microbiota

As Figure 4 shows, there are clear differences between in the profiles (Figures 4a and 4c) and the diversity (Figure 4b) of the frankfurter fed mices microbiota compared with all other groups. These differences exist on all phylogenetic levels (phylum, class, order, family, genus, and species). An analysis of covariance (ANCOVA) was used to compare the mean OTU levels of each phylum from post study between dietary groups, whilst controlling for baseline levels of each phylum. There was no significant difference in *Bacteroidetes* from the terminal sample between any groups, when controlling for baseline. The highest levels of *Firmicutes* were seen in the frankfurter consuming group, and the lowest levels were seen in the sausage consuming group, however when controlling for baseline, the there was no significant differences between any groups. The highest levels of *Actinobacteria* were seen in the frankfurter consuming group, and this was significantly higher than the control (p=0.002), sausage (0.001), and pork (<0.001) consuming groups. Similarly, the highest levels of *Proteobacteria* were seen in the frankfurter group and these were significantly higher than the pork (P=0.014), and sausage (P=0.011), but not the control (P=0.551). The control group had significantly higher levels of *Tenericutes* than the pork (P=0.001), sausage (P=0.001), and frankfurter (P=0.002) fed groups. The control group had significantly lower levels of *Verrucomicrobia* than the pork (P=0.551), sausage (P=0.551), and frankfurter groups (P=0.551).

**Figure 4.**
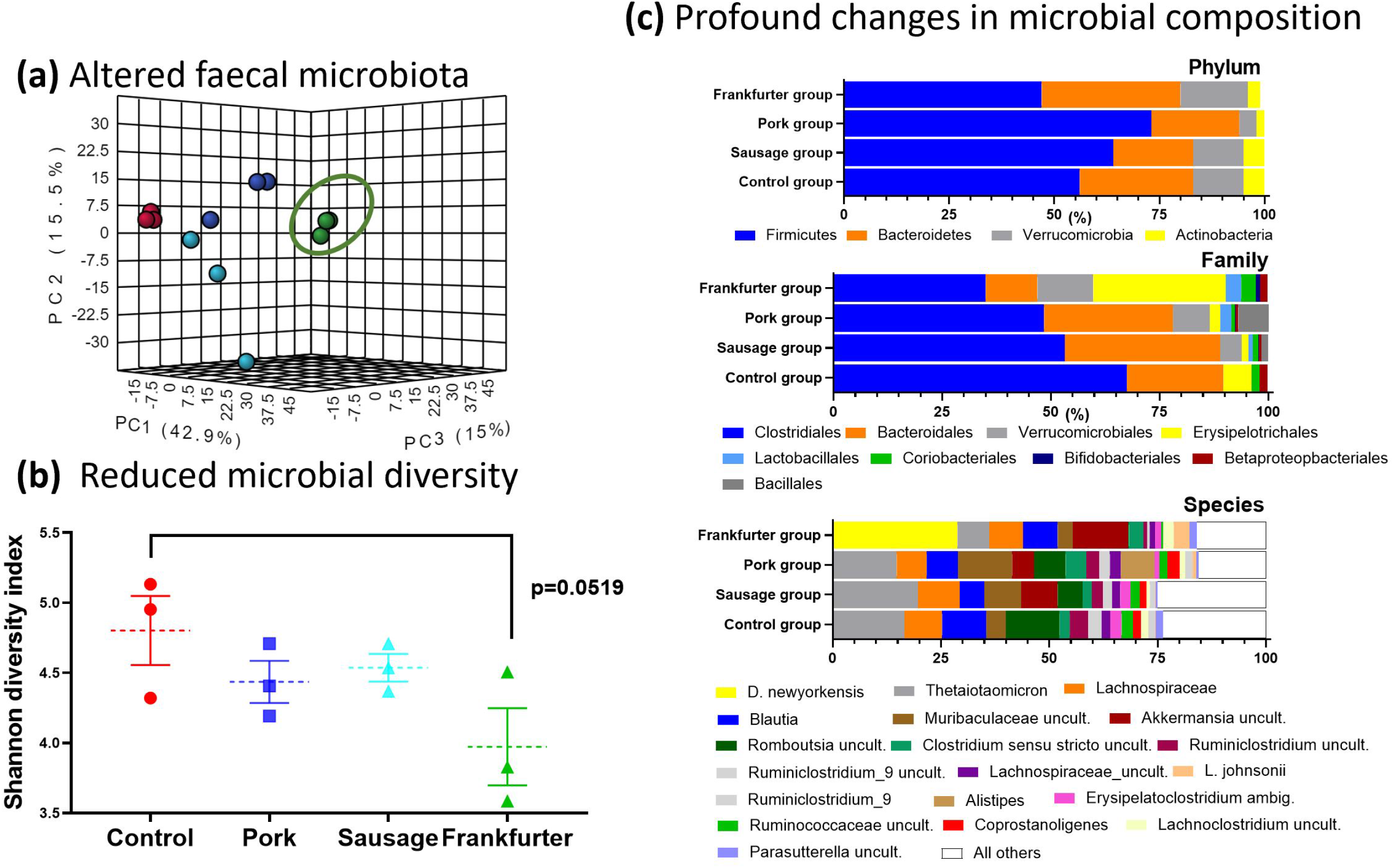
Faecal microbiome profiles differ most in frankfurter-fed mice with reduced micribial diversity and a profound shift in abundance of taxa between feeding groups. Faecal samples were collected at 56 days from each mouse cage for metagenomic sequencing providing cage replicates (n=3) for each treatment. **a)** shows scores plots of unsupervised PCA based on the first three principle components (PCs). Species level microbiome data explained 73.4% of the total variation (42.9%, 15.5% and 15.0% for PC1, PC2 and PC3, respectively). **b)** the Shannon diversity index as calculated by Qiime 2 indicates a reduction in microbial diversity in the frankfurter-fed animals. Individual values are shown along with means (dotted lines), and the error bars shown represent s.e.m. **c)** Relative abundance of microbial taxa is shown at the level of the phylum, the family, and the species. The taxa differed most profoundly in the frankfurter-fed animals. Bacterial species with an abundance of <1% were aggregated and referred to as ‘All others’ (white bars in figure). At the species level the dominance of *Dubiosiella Newyorkenesis* in the frankfurter group was a particularly striking observation. Control samples are shown in red, minced pork in dark blue, nitrite-free sausage in light blue, and frankfurter (nitrite-containing) in green. uncult. – uncultured bacteria; ambig. – ambiguous taxa.

The most abundant bacteria at the species level in the control, sausage and pork group was *Bacteriodes thetaiotaomicron*, the gut microbiome consisted of a mean (SD) relative abundance of 16.6 (9.37), 14.7 (9.07), and 19.7 (3.71)% respectively. The frankfurter group tended to have lower abundance than all 3 other groups at 7.3 (2.9)% (not significant). *Duboisiella newyorkenesis* was the most abundant species in the frankfurter group, making up a total of 28.8 (1.37)% of all species. This species was not detectable in any other group. All four groups displayed a high abundance of bacteria from the *Lachnospiraceae* family, which was largely attributed to higher levels of bacteria from the *Blautia* genus (see Supplementary Table 1). The sausage group was the only group with measureable *alistipes*, which was the 3^rd^ most abundant species 11.7 (5.5)%.

The most abundant bacteria at the family level in all four dietary groups is *Clostridiales*, the next most abundant in the control, sausage, and pork groups is *Bacteroidales*, whilst *Erysipelotrichales* is the second most abundant in the frankfurter group.

## Discussion

This study found that a modest inclusion of a sodium-nitrite containing pork frankfurter in the diet of APC^min^ mice significantly increased the number of intestinal tumours present. A similar inclusion of either nitrite-free pork sausage or minced pork did not increase tumour numbers. However, the numbers of intestinal foci (ACF and MDF) did not differ between any groups. Frankfurter consumption was also associated with higher levels of oxidative stress, as indicated by serum MDA and urinary DHN-MA, than any other dietary group. Unbiased PCA analysis of serum metabolomics data showed a distinct clustering of the frankfurter group which distinguished this diet from all others. This metabolic shift resulted from vastly more serum metabolites being significantly altered in Frankfurter-fed mice than any other group. When hierarchical clustering was performed on a shortlist of the 50 most affect metabolites across all groups it was demonstrated that the metabolic status of mice fed minced pork and those fed pork sausage were most similar to one another. It also illustrated the widerspread metabolic divergance of frankfurter-fed mice. Of the many metabolites profiled trimethylamine N-oxide (TMAO) stood out, since it has previously has linked to CRC development ^9^, and here it was uniquely elevated in the frankfurter group.

Intestinal microbiota play a significant role in TMAO production and precursor molecules of TMAO found in red meat, eggs, milk and fish. However, the increases in circulating TMAO here most likely result from alterations in the gut microbiome. It is clear that all dietary interventions in this study impacted the composition of faecal microbiota to some extent. However, the reduced microbial diversity (Shannon index) of the frankfurter group suggested that this diet resulted in gut dysbiosis. The most remarkable observation was the sheer dominance of *D. newyorkenesis* in the frankfurter group. The levels of this organism in all other groups fell well below threshold cut-off values. *D. newyorkenesis* is a poorly characterised organism and we questioned whether *D. newyorkenesis* is a driver of TMAO formation which is subsequently detected in the blood circulation. The genome of this recently identified bactierial species ^10^ has was sequenced recently (but is not yet fully annotated). We utilized the makeblastdb application and produced a searchable BLAST databases from FASTA files obtained from D. newyorkenesis and carried out pairwise alignments to search for gene sequences encoding enzmyes involved in TMA-synthesis pathways (including choline TMA-lyase (cutC) and carnitine oxygenase (cntA)) ^11^ but we did not find any significant alignments, which would suggest that this organism is not involved in TMAO production.

Nonetheless, at the genus level *Dubosiella* is positively associated with obesity in mice ^12^ and at the ‘family’ level, *Erysipeltrchaceae*, is positively associated with CRC in murine models and humans ^13,14^. It is not however clear if predominance of *Erysipeltrchaceae* is causative of CRC or if colonisation occurs in response to CRC. The *Clostridiales* family, although present in all four groups, was noticeably less prevalent in the frankfurter group. It may be that *Erysipeltrchaceae* to some extent displaced *Clostridiales*, but another explanation could be the known ability of nitrites to prevent the growth of organisms from the *clostridium* genus.

An in-depth literature search reveals that ten pre-clinical studies have thusfar evaluated the effect of nitrite consumption on the development of CRC. The deficiency of most of these studies is that cancer was chemically induced and not spontaneous. None-the-less, five studies reported nitrite exposure to increase cancer risk ^15-19^. Three studies reported that nitrite treatment made no difference ^20-22^, and one study reported nitrite to actually be beneficial ^23^. Our study has several advantages over previous studies in this area. Firstly, it is the only study to provide a nitrite-containing meat matrix to Apc^Min+/+^ mice, which could be described as a ‘gold-standard’ model of spontaneous CRC development. This minimises the known unpredictability of cancer development (especially in location and timing) of chemically-induced CRC models. Secondly, all previous pre-clinical studies linking nitrite to CRC development, provided a minimium of 50% processed meat in the diet. Not only are such levels unrealistic for humans, they disproportionately alter the overall nutritional profile of the diet. Our study has detected a measurable effect with just 15% dietary inclusion, and although this still represents a relatively high intake of processed meat it clearly demonstrates that lower dietary quantities can exacerbate the disease. Thirdly, this study has investigated other relatively unexplored mechanisms through which diets impacts CRC. The application of both metabolome and microbiome data makes this study particularly unique. Lipid peroxidation (i.e. the oxidative destruction of membrane lipids and production of compounds impairing cells and tissues) was evident and this is in keeping with other studies ^16,18^. The aformentioned rise in TMAO in the frankfurter group could also be linked linked to oxidative stress ^24^. TMAO is generated from the metabolism of a number of nutrients including carnitine and choline which are present in pork meat. Increased TMAO in the circulation apparently induces superoxide production, a reactive oxygen species linked to oxidative stress ^24^. TMAO’s production of reactive oxygen species has been confirmed in cells *in vitro* ^25^. The association between TMAO, oxidative stress and cancer still needs to be verified, but mounting evidence suggests that TMAO could play a role in chronic inflammation as a cause of carcinogenesis in CRC ^26^. Furthermore, *in silico* studies suggest that TMAO metabolism and CRC share many similar genetic pathways ^27^.

In sharp contrast to the frankfurter group, the minced pork and pork sausage group did not display elevated serum TMAO. The TMAO responses did not mirror those of common TMAO precusors. Choline levels were the same in all groups. As would be expected carnitine (and several acylcarnitines) were high in all three pork diet interventions. Another TMAO precursor betaine (found typically in plants but not in meat) did not differ in any groups either. Therefore, in these experiments the supply of precursors does not appear to be important factor in TMAO production, and the suspicion would be that altered microbiota are more pivotal. A limitation of this study is the fact that the microbial profiling undertaken involved the analysis of 16S rRNA amplicon rather than a whole-metagenomics approach, thus it was not possible to evaluate whether there was an abundance of bacterial species with genes encoding the enzymatic machinery involved in TMAO production.

An obvious advantage of the present study design is that it reflects real-world human food choices. To this end it directly compared commercially available pork products on an equivalent gram-for-gram basis. The unfortunate consequence of this approach is that the nutritional properties of the pelleted diets cannot be precisely-matched, or be isocaloric, which is a potential limitation. Another limitation is that is was not possible to record food intake, and we cannot pinpoint the underlying reasons for the higher bodyweights of mice in the frankfurter group. It is possible that mice found this pelleted diet more palatable and consumed more, but there are also differences in the levels of fibre, and the types of fat in the diets which could have contributed also to this effect. It is not known if raised bodyweight exacerbated CRC pathology in this group, but this is very questionable given that the pork sausage group had the lowest body weight, but also the second highest number of tumours overall. There were metabolic differences evident in the frankfurter group, for instance, the significantly elevated diglycerides and triglycerides, which were likely explained by the higher saturated fat content of this diet. The decreased phosphatidylcholine and lysophosphatidylcholine (lysoPC) levels may also be a consequence of this, but equally, similar observations have been noted in CRC patients ^28^. Increased lysoPC metabolism in tumour cells *in vitro* is one explanation for a decline in their blood concentrations.

The results from this study clearly show that not all processed meats carry the same risk of CRC development, and that the consumption of nitrite-containing processed meat exacerbates CRC pathology in a genetically predisposed murine model.

However, higher dietary fat and increased body weight cannot completely be ruled out as potential confounding factors. The metabolomic and gut microbiome data demonstrate the systemic alterations that can occur as result of simple dietary choices. These also provide important mechanistic data. Further dose-response studies should be performed, perhaps using humanised or colon-specific CRC mouse models, and employing frankfurter as a model food matrix. Such work is required to determine the levels at which nitrite inclusion becomes harmful.

## Methods

### Animal models

Forty female Apc^Min/+^ mice were purchased from Charles River and delivered to Axis Bioservices (Coleraine, Northern Ireland) where the study was conducted. All mice were four weeks old on delivery. The mice were all fed an ad lib maintenance diet of AIN76 for a two week acclimitization period (Figure 1a). Mice were caged in groups of 3 or 4 in a 12 hour light, 12 hour dark cycle. Mice had unlimited access to their respective diets and water for the duration of the study which was 12 weeks, with the exception of 12 hours prior to blood collection, when they were fasted. Body weight was measured every 3–4 days. We attempted to measure food intake in a similar manner, but since diets were required (for welfare reasons) to be placed in dishes in home cages this led to extremely erroneous data and was subsequently abandoned. Mice were randomly assigned to one of four treatment groups, described below. Studies in mice underwent Ethical Review and were performed in accordance with the UK Animals (Scientific Procedures) Act 1986.

### Diets

Each treatment group received one of four diets for the duration of the study, the dietary groups included; Control, which consisted of 100% AIN76 (Altromin, Germany). Sausage meat (15%)/AIN76 (85%), Frankfurter (15%)/AIN76 (85%), unprocessed pork (15%)/AIN76 (85%). Sausage meat was manufactured and provided by Finnebrogue Artisan (Downpatrick, UK) as was the unprocessed pork. Nitrite-containing Frankfurter was manufactured by Herta (Illkirch, France) and purchased from a local supermarket. All meat was freeze dried by European Freeze Dry (UK), and powderized. Altromin then incorporated this into pellet composed of 15% meat powder and 85% AIN76. The nutritional properties of each diet were analysed after pelleting (Supplementary Table 2).

### Biological samples

Blood samples were collected from the tails of all animals at 3 time points, week 0, week 3, week 6, a final cardiac puncture blood was taken at the termination of the study (week 8). Urine was collected from individual animals at baseline and week 8. Faecal samples at the same timepoints but were collected from each cage.

### Tumor scoring in the gastrointestinal tract

Animals were sacrificed using CO_2_ asphyxiation and their organs harvested. As described previously^29^, the intestinal tract was removed washed in phosphate buffered saline and sectioned into the duodenum, jejunum, ileum, and colon. The sections were opened longitudinally and fixed in 10% formalin, colons were stained for 5 minutes in 0.05% methylene blue solution (Sigma), and small intestine were stained for 48 hours in 0.05% methylene blue. Using x25 magnification all tumors present were counted by one researcher in a blinded manner. The minimum size for accepting and counting a tumor was 0.5mm diameter.

### Identification of aberrant crypt foci and mucin depleted foci

Aberrant crypt foci (ACF) were measured in the colon by a well described method ^30^. The colons were washed in saline, and stained with 0.2% methylene blue for 5 mins. They were placed on a glass slide and visualised using a magnification of 32x on a light microscope. Mucin depleted foci (MDF) were also measured in the colon, the sections were washed in saline, and stained with 1% high-iron diamine alcian blue procedure for 30 mins. All scoring was completed by one researcher and verified by a second researcher in a blinded manner.

### Lipid peroxidation

Serum samples from every time point were analysed for malondialdehyde (MDA) using ELISA methodologies (Abcam, Cambridge UK). In brief, thiobarbituric acid, serum and standards were added to 96-well microtiter plate and incubated at 95°C for 60 mins and cooled on ice. An aliquot from each well was added to the colorimetric plate and measured at OD532 nm using a plate reader. Urine samples from baseline and week 8 were analysed for 1,4-dihydroxynonane mercapturic acid (DHN-MA) using a Bertin Bioreagent enzyme immunoassay (Wolverhampton UK). Briefly, buffer, standards and urine were added to each well, the detection antibody was then added, and the plate was incubated overnight at 4°C. Ellman’s reagent was added, and the absorbance was measured at 414nm.

### Metabolite profiling

The MxP Quant 500 kit (Biocrates, Life science AG, Innsbruck, Austria) was used to quantify amino acids, acylcarnitines, bile acids, biogenic amines, carboxylic acids, ceramides, cholesterol esters, diacylglycerols, fatty acids, glycerophospholipids, glycosylceramides, sphingolipids, and triacylglycerols in serum samples from baseline and week 8. Serum (10μL) samples were analysed using a triple-quadrupole mass spectrometer (Xevo TQ-S, Waters Corporation, Milford, CT, USA). Samples were dispensed into a 96-well plate, dried at room temperature for 30 min and then 50 μL of phenylisothiocyanate (PITC) was added. After a further 20 min incubation at room temperature samples were dried under nitrogen for 1 h. Ammonium acetate (3 mM) was added, and the plate was shaken at room temperature for 30 min and centrifuged for 2 min at 500 *g*. Each well then had 150 μL transferred from it to a 96 deep well plate along with and equal volume of dH_2_O. A 96-well plate containing 490 μL of FIA solvent is shaken along with 10 μL of extract at room temperature for 5 mins. From each well 5 μL of sample Samples were injected through the MxP Quant 500 kit column system for amino acids and biogenic amines analysis. Other metabolites were semi-quantified using the same mass spectrometer without column separation by the flow injection analysis (FIA). Metabolite concentrations were calculated using the Analyst/MetIDQ software and expressed as μM in serum.

### Microbial profiling

A QIAamp DNA stool mini kit (Qiagen, Manchester, UK) was used to extract DNA from faecal samples. All reagents, unless otherwise stated were provided by Qiagen. Faecal samples were weighed (220mg) and added to a 2ml Eppendorf tube along with 1.4 ml of ASL buffer, which was vortexed and heated for 5 mins at 95°C. The contents were added to a 2 ml screw capped tube containing 0.5 g of both 0.1 mm and 0.5 mm zirconium beads. The tube was then added to a FastPrep 24 g5 bead beater (MP biomedicals, Eschwege, Germany) for 5 × 20 s bursts. The suspension was subsequently spun at 14,000 *g* for 3 mins. The supernatant (1.2 ml) was collected and transferred into a QIAamp silica-gel membrane tube and spun for 3 min, and DNA extracted in accordance with the manufacturer’s instructions. Amplicon sequencing was carried out on an Illumina MiSeq. Variable regions V4-V5 of the bacterial 16S ribosomal RNA gene were amplified from extracted DNA using PCR conditions and custom primers as described in the Microbiome Helper protocol ^31^. The forward (515FB = GTGYCAGCMGCCGCGGTAA) and reverse (926R = CCGYCAATTYMTTTRAGTTT) primers used Nextera Illumina index tags and sequencing adapters fused to the 16S sequences. Each sample was amplified with a different combination of index barcodes to allow for sample identification after multiplex sequencing. After amplification, paired-end 300 + 300 bp v3 sequencing was performed for all samples on an Illumina MiSeq. Raw sequencing reads were trimmed as previously described ^32^.

### Analysis of 16S sequencing data

Analysis of 16S sequencing data was carried out on the Hardiman Laboratory GRANDE server. Primer sequences were removed from the sequencing reads using cutadapt (v 1.14) ^33^, and primer-trimmed files were imported into QIIME2 (v. 2019.10.0) ^34^. Forward and reverse paired-end reads were joined using VSEARCH (v 2.9.0) ^35^ and inputted to Deblur ^36^ to correct reads and obtain amplicon sequence variants (ASVs). Taxonomy was assigned to ASVs using the SILVA rRNA gene database ^37^ and the “feature-classifier” option in QIIME2, sklearn. Rare sequences were removed from the feature table. Any sequences below an arbitrary cut-off frequency of 100 were also removed. Estimates of alpha-diversity (Observed ASVs, Shannon Diversity), beta-diversity (weighted UniFrac), and relative abundance of ASVs were obtained using QIIME2 (v 2019.10.0). Differential abundance tests were done using the Analysis of compositions of microbiomes (ANCOM)plugin ^38^.

### Statistical analysis

All data were tested for normality using Kologorov-Smirnoff and where appropriate data was transformed. Group comparisons for tumour numbers, ACF and MDF were performed by a One-way ANOVA with LSD post-hoc test (two-tailed; SPSS version 21). Lipid peroixdation markers DHN-MA and MDA at week 8 were compared between groups whilst controlling for baseline concentrations by an ANCOVA. Metabolomic data were cube-root transformed, underwent mean-centered scaling and compared using univariate (ANOVA), multivariate and hierarchial clustering tools within MetaboAnalyst 4.0 (http://metaboanalyst.ca).

## Data Availability Statement

Raw mass spectrometry data is available on the Metabolomics Workbench (https://www.metabolomicsworkbench.org/) under study ST002313, [http://dx.doi.org/doi: 10.21228/M81M72]. The 16S rRNA Seq data have been submitted to the European Nucleotide Archive (ENA), accession number PRJEB56215.

## Acknowledgements

Studies investigating links between meat consumption and CRC are in-part supported by Agri-Food QUEST, a membership-based, industry-led Innovation Centre for agri-food business in Northern Ireland. Agri-Food QUEST is supported by funding from Invest Northern Ireland (InvestNI) but also were partially supported by commercial funding from: Finnebrogue Artisan, Karro Food group, and Cranswick. These industrial partners were not involved design or writing of the manuscript, the analysis/interpretation of the data, or the decision to publish.

## Author Contributions

W.C. organised, managed and conducted laboratory work. X.P. conducted metabolomic studies. G.H., J.M. and A.H. performed metagomic sequencing analysis. W.C. and B.D.G. wrote the manuscript. B.D.G, G.H. and C.T.E edited the manuscript. All authors approved the final text of the manuscript.

