## Supplementary Material for "Dietary inclusion of nitrite-containing frankfurter exacerbates colorectal cancer pathology, alters metabolism and causes gut dybiosis in APC^min^ mice"

### Supplementary Table 1: Percentage Species Abunandance in Individual Faecal Samples

| | **GROUP** | | **Control** | **Control** | **Control** | **Pork** | **Pork** | **Pork** | **Sausage** | **Sausage** | **Sausage** | **Frankfurter** | **Frankfurter** | **Frankfurter** | **MEAN (all samples)** |
|---|---|---|---|---|---|---|---|---|---|---|---|---|---|---|---|
| | **Species level output from Qiime 2** | **Label in Figure 4C** | PN13 | PN14 | PN15 | PN16 | PN17 | PN18 | PN19 | PN20 | PN21 | PN22 | PN23 | PN24 | |
| 1 | D_0__Bacteria;D_1__Bacteroidetes;D_2__Bacteroidia;D_3__Bacteroidales;D_4__Bacteroidaceae;D_5__Bacteroides;D_6__Bacteroides_thetaiotaomicron | Thetaiotaomicron | 17.23529 | 25.57786 | 6.859517 | 5.011826 | 16.23198 | 22.94605 | 16.06236 | 19.40043 | 23.53271 | 5.6274155 | 5.6095688 | 10.661665 | 14.56305568 |
| 2 | D_0__Bacteria;D_1__Firmicutes;D_2__Clostridia;D_3__Clostridiales;D_4__Lachnospiraceae;__;__ | Lachnospiraceae | 7.42517 | 12.77909 | 5.840861 | 5.140462 | 7.285231 | 8.387702 | 6.226257 | 10.45234 | 12.18304 | 10.822131 | 8.0882531 | 4.4439867 | 8.256210614 |
| 3 | D_0__Bacteria;D_1__Firmicutes;D_2__Clostridia;D_3__Clostridiales;D_4__Lachnospiraceae;D_5__Blautia;__ | Blautia | 11.85878 | 9.658089 | 9.073173 | 7.977924 | 5.801706 | 7.68189 | 3.529165 | 5.502314 | 7.949205 | 9.8827838 | 6.9705419 | 7.2259505 | 7.759293196 |
| 4 | D_0__Bacteria;D_1__Bacteroidetes;D_2__Bacteroidia;D_3__Bacteroidales;D_4__Muribaculaceae;D_5__uncultured_bacterium;D_6__uncultured_bacterium | Muribaculaceae uncult. | 3.910041 | 3.926034 | 5.505998 | 14.09021 | 9.230656 | 14.31353 | 9.488149 | 11.58105 | 4.489316 | 3.1276899 | 3.1987716 | 3.9918905 | 7.237778308 |
| 5 | D_0__Bacteria;D_1__Firmicutes;D_2__Erysipelotrichia;D_3__Erysipelotrichales;D_4__Erysipelotrichaceae;D_5__Dubosiella;D_6__Dubosiella_newyorkensis | D. newyorkensis | 0.016285 | 0.019672 | 0.015008 | 0.020748 | 0.069873 | 0.070321 | 0.008405 | 0.024136 | 0.01981 | 27.553019 | 30.25094 | 28.628425 | 7.224720019 |
| 6 | D_0__Bacteria;D_1__Verrucomicrobia;D_2__Verrucomicrobiae;D_3__Verrucomicrobiales;D_4__Akkermansiaceae;D_5__Akkermansia;D_6__uncultured_bacterium | Akkermansia uncult. | 0.083054 | 0.052962 | 0.065659 | 7.655919 | 2.382318 | 5.053327 | 7.291141 | 11.7713 | 6.175884 | 9.8833615 | 12.346547 | 16.264623 | 6.585508329 |
| 7 | D_0__Bacteria;D_1__Firmicutes;D_2__Clostridia;D_3__Clostridiales;D_4__Peptostreptococcaceae;D_5__Romboutsia;D_6__uncultured_bacterium | Romboutsia uncult. | 13.43843 | 6.60064 | 16.98136 | 11.95485 | 6.201824 | 3.748486 | 4.840309 | 5.803305 | 6.630144 | 0.0831894 | 0.0800097 | 0.4282446 | 6.399232794 |
| 8 | D_0__Bacteria;D_1__Firmicutes;D_2__Clostridia;D_3__Clostridiales;D_4__Clostridiaceae_1;D_5__Clostridium_sensu_stricto_1;D_6__uncultured_bacterium | Clostridium sensu stricto uncult. | 2.187897 | 1.482193 | 3.610322 | 6.646749 | 7.604442 | 0.01172 | 2.161708 | 2.342619 | 1.758976 | 2.8994968 | 3.2270578 | 3.6677255 | 3.133408818 |
| 9 | D_0__Bacteria;D_1__Firmicutes;D_2__Clostridia;D_3__Clostridiales;D_4__Ruminococcaceae;D_5__Ruminiclostridium;D_6__uncultured_bacterium | Ruminiclostridium uncult. | 4.256913 | 5.431682 | 3.06066 | 1.324536 | 2.806708 | 4.958263 | 1.507816 | 2.64432 | 3.532297 | 1.0109822 | 0.691801 | 1.1307827 | 2.696396755 |
| 10 | D_0__Bacteria;D_1__Firmicutes;D_2__Clostridia;D_3__Clostridiales;D_4__Ruminococcaceae;D_5__Ruminiclostridium_9;__ | Ruminiclostridium_9 | 3.295281 | 3.082417 | 2.6592 | 2.596788 | 1.841718 | 2.846688 | 1.698605 | 2.326291 | 2.39494 | 0.575971 | 0.5115772 | 0.6147208 | 2.037016404 |
| 11 | D_0__Bacteria;D_1__Firmicutes;D_2__Clostridia;D_3__Clostridiales;D_4__Lachnospiraceae;D_5__uncultured;D_6__uncultured_bacterium | Lachnospiraceae uncult. | 1.828812 | 2.288736 | 2.072019 | 1.75028 | 1.711533 | 4.090974 | 1.223735 | 1.696624 | 2.925706 | 1.3639594 | 1.1581202 | 1.1708967 | 1.940116273 |
| 12 | D_0__Bacteria;D_1__Bacteroidetes;D_2__Bacteroidia;D_3__Bacteroidales;D_4__Rikenellaceae;D_5__Alistipes;__ | Alistipes | 0 | 0 | 0.006566 | 7.788705 | 15.57223 | 0.004558 | 0.002521 | 0.003549 | 0 | 0.0132872 | 0.0121227 | 0.0108416 | 1.951198181 |
| 13 | D_0__Bacteria;D_1__Firmicutes;D_2__Erysipelotrichia;D_3__Erysipelotrichales;D_4__Erysipelotrichaceae;D_5__Erysipelatoclostridium;Ambiguous_taxa | Erysipelatoclostridium ambig. | 5.168876 | 1.69253 | 0.983951 | 1.999253 | 0.78479 | 0.509174 | 1.559086 | 3.140529 | 2.123067 | 1.9139336 | 1.8685093 | 0.6982014 | 1.870158287 |
| 14 | D_0__Bacteria;D_1__Firmicutes;D_2__Clostridia;D_3__Clostridiales;D_4__Ruminococcaceae;D_5__uncultured;__ | Ruminococcaceae uncult. | 2.34749 | 2.53766 | 2.857116 | 1.088012 | 1.074581 | 3.443112 | 1.713733 | 2.433484 | 2.273348 | 0.3050278 | 0.2957934 | 0.4586012 | 1.735663257 |
| 15 | D_0__Bacteria;D_1__Firmicutes;D_2__Clostridia;D_3__Clostridiales;D_4__Ruminococcaceae;D_5__[Eubacterium]_coprostanoligenes_group;__ | Coprostanoligenes | 3.330293 | 2.418873 | 0.006566 | 4.489813 | 1.533539 | 2.414345 | 0.339553 | 0.434449 | 3.938057 | 0.0600812 | 0.0509153 | 0.0205991 | 1.586423684 |
| 16 | D_0__Bacteria;D_1__Firmicutes;D_2__Clostridia;D_3__Clostridiales;D_4__Lachnospiraceae;D_5__Lachnoclostridium;D_6__uncultured_bacterium | Lachnoclostridium uncult. | 1.516138 | 1.643351 | 1.665869 | 1.389269 | 1.33348 | 1.157688 | 0.519415 | 0.741119 | 1.213181 | 2.9959734 | 2.4334263 | 1.4256752 | 1.502882178 |
| 17 | D_0__Bacteria;D_1__Firmicutes;D_2__Clostridia;D_3__Clostridiales;D_4__Ruminococcaceae;D_5__Ruminiclostridium_9;D_6__uncultured_bacterium | Ruminiclostridium_9 uncult. | 1.33456 | 1.183333 | 2.515688 | 2.165235 | 1.328332 | 1.701371 | 1.022021 | 1.426868 | 1.234357 | 0.1322942 | 0.1026387 | 0.1550354 | 1.191811107 |
| 18 | D_0__Bacteria;D_1__Firmicutes;D_2__Bacilli;D_3__Lactobacillales;D_4__Lactobacillaceae;D_5__Lactobacillus;D_6__Lactobacillus_johnsonii | L. johnsonii | 0.045598 | 0.046153 | 0.038458 | 1.543633 | 0.700206 | 0.253285 | 0.090772 | 0.098674 | 0.033472 | 3.6568669 | 3.973815 | 3.0866138 | 1.130628757 |
| 19 | D_0__Bacteria;D_1__Proteobacteria;D_2__Gammaproteobacteria;D_3__Betaproteobacteriales;D_4__Burkholderiaceae;D_5__Parasutterella;D_6__uncultured_bacterium | Parasutterella uncult. | 1.606521 | 0.976023 | 2.48192 | 0.498776 | 0.458223 | 0.533266 | 0.49336 | 0.598433 | 0.413957 | 1.1091918 | 1.171051 | 2.8459295 | 1.098887555 |
| 20 | D_0__Bacteria;D_1__Firmicutes;D_2__Clostridia;D_3__Clostridiales;D_4__Lachnospiraceae;D_5__uncultured;__ | All others | 1.17171 | 1.384591 | 1.767172 | 0.880534 | 1.171668 | 0.997513 | 0.751387 | 1.443905 | 1.258265 | 0.5909913 | 0.2966016 | 0.2233377 | 0.994806326 |
| 21 | D_0__Bacteria;D_1__Bacteroidetes;D_2__Bacteroidia;D_3__Bacteroidales;D_4__Muribaculaceae;__;__ | All others | 0.907892 | 1.075895 | 0.708182 | 0.465579 | 0.166961 | 0.973421 | 1.138847 | 1.426868 | 0.822449 | 1.0161815 | 1.2243908 | 1.0819953 | 0.917388525 |
| 22 | D_0__Bacteria;D_1__Firmicutes;D_2__Clostridia;D_3__Clostridiales;D_4__Ruminococcaceae;D_5__Intestinimonas;D_6__uncultured_bacterium | All others | 0.710028 | 0.759633 | 4.177805 | 0.114528 | 0.003678 | 0.003907 | 0.928728 | 1.379305 | 1.129843 | 0.9896071 | 0.5277407 | 0.7144638 | 0.95327223 |
| 23 | D_0__Bacteria;D_1__Actinobacteria;D_2__Coriobacteriia;D_3__Coriobacteriales;D_4__Atopobiaceae;D_5__Coriobacteriaceae_UCG-002;D_6__uncultured_bacterium | All others | 0.001629 | 0.003026 | 0.001876 | 0.005809 | 0.008826 | 0.018882 | 0.009245 | 0.018457 | 0.018444 | 2.9474463 | 3.3127248 | 3.0399948 | 0.782196711 |
| 24 | D_0__Bacteria;D_1__Firmicutes;D_2__Clostridia;D_3__Clostridiales;D_4__Lachnospiraceae;D_5__Lachnospiraceae_UCG-006;__ | All others | 0.846823 | 1.035795 | 0.754144 | 0.36848 | 0.472198 | 0.422576 | 0.854766 | 0.989579 | 1.558828 | 0.5343763 | 0.4073221 | 0.1420254 | 0.698909274 |
| 25 | D_0__Bacteria;D_1__Firmicutes;D_2__Clostridia;D_3__Clostridiales;D_4__Ruminococcaceae;D_5__Ruminococcaceae_UCG-014;__ | All others | 0.225548 | 0.291294 | 0.284211 | 0.029877 | 0.024272 | 0.01172 | 0.103379 | 0.181021 | 0.017077 | 2.721564 | 3.5858892 | 0.4564329 | 0.661023666 |
| 26 | D_0__Bacteria;D_1__Firmicutes;D_2__Clostridia;D_3__Clostridiales;D_4__Ruminococcaceae;__;__ | All others | 1.205908 | 1.120535 | 1.711831 | 0.229055 | 0.326567 | 0.681721 | 0.115145 | 0.30596 | 0.743893 | 0.5823257 | 0.4719764 | 0.3686156 | 0.655294409 |
| 27 | D_0__Bacteria;D_1__Firmicutes;D_2__Bacilli;D_3__Bacillales;D_4__Planococcaceae;D_5__Sporosarcina;D_6__Sporosarcina_sp._106 | All others | 0 | 0.00227 | 0.002814 | 0.00166 | 0 | 0 | 7.978652 | 0.06176 | 0.01093 | 0.0127095 | 0.0064654 | 0.0032525 | 0.673376033 |
| 28 | D_0__Bacteria;D_1__Firmicutes;D_2__Erysipelotrichia;D_3__Erysipelotrichales;D_4__Erysipelotrichaceae;D_5__Faecalibaculum;D_6__uncultured_bacterium | All others | 0 | 0 | 9.002823 | 0 | 0 | 0 | 0 | 0.00142 | 0 | 0 | 0.0072736 | 0 | 0.750959727 |
| 29 | D_0__Bacteria;D_1__Firmicutes;D_2__Clostridia;D_3__Clostridiales;D_4__Christensenellaceae;D_5__uncultured;D_6__uncultured_bacterium | All others | 0.606618 | 0.259516 | 0.642523 | 1.097971 | 0.600177 | 0.773528 | 0.732056 | 0.856121 | 1.372343 | 0.1149631 | 0.0759688 | 0.1225105 | 0.604524535 |
| 30 | D_0__Bacteria;D_1__Firmicutes;D_2__Clostridia;D_3__Clostridiales;D_4__Ruminococcaceae;D_5__Oscillibacter;D_6__uncultured_bacterium | All others | 0.661173 | 0.821675 | 1.816886 | 0.352712 | 0.231686 | 0.319048 | 0.761473 | 1.149303 | 0.463823 | 0.2709432 | 0.153554 | 0.3100708 | 0.609362168 |
| 31 | D_0__Bacteria;D_1__Firmicutes;D_2__Bacilli;D_3__Lactobacillales;D_4__Carnobacteriaceae;D_5__Atopostipes;D_6__uncultured_bacterium | All others | 0 | 0.001513 | 0 | 0 | 0.015446 | 0 | 7.171794 | 0.026266 | 0.002049 | 0.0040439 | 0 | 0.0021683 | 0.601939982 |
| 32 | D_0__Bacteria;D_1__Firmicutes;D_2__Clostridia;D_3__Clostridiales;D_4__Lachnospiraceae;D_5__Tyzzerella;D_6__uncultured_bacterium | All others | 0.689672 | 0.62874 | 0.223242 | 0.209137 | 0.522948 | 0.987095 | 0.557237 | 0.653804 | 0.993907 | 0.2120174 | 0.1567867 | 0.1832237 | 0.501484054 |
| 33 | D_0__Bacteria;D_1__Firmicutes;D_2__Bacilli;D_3__Bacillales;D_4__Staphylococcaceae;D_5__Staphylococcus;D_6__Staphylococcus_lentus | All others | 0.013842 | 0.012862 | 0.00469 | 0.00249 | 0.275816 | 0.009767 | 5.729534 | 0.025556 | 0.013662 | 0.0046216 | 0 | 0.0195149 | 0.509363018 |
| 34 | D_0__Bacteria;D_1__Firmicutes;D_2__Clostridia;D_3__Clostridiales;D_4__Lachnospiraceae;D_5__Lachnospiraceae_NK4A136_group;__ | All others | 0.238576 | 0.165697 | 0.826369 | 0.151044 | 0.094881 | 0.225287 | 0.085729 | 0.149076 | 0.341549 | 1.2877024 | 1.0603305 | 0.2526101 | 0.406570771 |
| 35 | D_0__Bacteria;D_1__Firmicutes;D_2__Clostridia;D_3__Clostridiales;D_4__Lachnospiraceae;D_5__Lachnospiraceae_UCG-006;D_6__uncultured_bacterium | All others | 0.061069 | 0.113491 | 0.941741 | 0.580107 | 0.466314 | 0.496152 | 0.263069 | 0.332936 | 0.562872 | 0.4974032 | 0.456621 | 0.0823964 | 0.404514352 |
| 36 | D_0__Bacteria;D_1__Firmicutes;D_2__Clostridia;D_3__Clostridiales;D_4__Ruminococcaceae;D_5__Oscillibacter;D_6__unidentified | All others | 0.845194 | 0.901119 | 0.083481 | 0.478028 | 0.441306 | 0.69865 | 0.17566 | 0.293182 | 0.212443 | 0.3148487 | 0.1147614 | 0.1940653 | 0.396061592 |
| 37 | D_0__Bacteria;D_1__Firmicutes;D_2__Bacilli;D_3__Bacillales;D_4__Staphylococcaceae;D_5__Staphylococcus;__ | All others | 0.004886 | 0.009836 | 0.005628 | 0.00415 | 4.450574 | 0.007162 | 0.007564 | 0.004259 | 0.002732 | 0.0034662 | 0.0129309 | 0.0054208 | 0.376550732 |
| 38 | D_0__Bacteria;D_1__Firmicutes;D_2__Clostridia;D_3__Clostridiales;D_4__Ruminococcaceae;D_5__Intestinimonas;__ | All others | 0.364785 | 0.684729 | 0.040334 | 0.253123 | 0.529568 | 0.914821 | 0.181543 | 0.135588 | 0.756189 | 0.0808786 | 0.0412171 | 0.0628815 | 0.337138003 |
| 39 | D_0__Bacteria;D_1__Actinobacteria;D_2__Coriobacteriia;D_3__Coriobacteriales;D_4__Eggerthellaceae;D_5__Enterorhabdus;Ambiguous_taxa | All others | 0.961632 | 0.50617 | 0.146326 | 1.179302 | 0.904678 | 0.108086 | 0 | 0 | 0.144817 | 0.0017331 | 0 | 0 | 0.329395346 |
| 40 | D_0__Bacteria;D_1__Firmicutes;D_2__Clostridia;D_3__Clostridiales;D_4__Lachnospiraceae;D_5__Lachnoclostridium;__ | All others | 0.25649 | 0.161914 | 0.176342 | 0.242334 | 0.141954 | 1.122527 | 0.208438 | 0.352813 | 0.324471 | 0.0935881 | 0.2238655 | 0.0162625 | 0.276749868 |
| 41 | D_0__Bacteria;D_1__Firmicutes;D_2__Erysipelotrichia;D_3__Erysipelotrichales;D_4__Erysipelotrichaceae;D_5__Turicibacter;__ | All others | 0.660359 | 0.500874 | 0.9333 | 0.759368 | 0.16549 | 0.008465 | 0.121029 | 0.18457 | 0.006148 | 0.0612366 | 0.1551703 | 0.119258 | 0.306272158 |
| 42 | D_0__Bacteria;D_1__Firmicutes;D_2__Clostridia;D_3__Clostridiales;D_4__Clostridiales_vadinBB60_group;D_5__uncultured_bacterium;D_6__uncultured_bacterium | All others | 0.054555 | 0.354849 | 0.033768 | 0.00166 | 0.008826 | 0.024091 | 0.266431 | 0.434449 | 0.006148 | 0.6528056 | 0.6045177 | 1.0635645 | 0.292138743 |
| 43 | D_0__Bacteria;D_1__Firmicutes;D_2__Clostridia;D_3__Clostridiales;D_4__Ruminococcaceae;D_5__Ruminiclostridium_5;D_6__uncultured_bacterium | All others | 0.349314 | 0.282214 | 0.473685 | 0.3718 | 0.318476 | 0.464247 | 0.189948 | 0.340035 | 0.359992 | 0.0502603 | 0.076777 | 0.0617973 | 0.278212149 |
| 44 | D_0__Bacteria;D_1__Actinobacteria;D_2__Coriobacteriia;D_3__Coriobacteriales;D_4__Eggerthellaceae;D_5__Parvibacter;D_6__uncultured_bacterium | All others | 0.531707 | 0.335177 | 0.573112 | 0.239844 | 0.088997 | 0.223333 | 0.353001 | 0.345714 | 0.409175 | 0.056615 | 0.0872833 | 0.1431096 | 0.282255602 |
| 45 | D_0__Bacteria;D_1__Firmicutes;D_2__Clostridia;D_3__Clostridiales;D_4__Peptococcaceae;D_5__uncultured;D_6__uncultured_organism | All others | 0 | 0 | 0.001876 | 1.552762 | 0.630332 | 0.774179 | 0 | 0 | 0 | 0.0034662 | 0 | 0 | 0.24688462 |
| 46 | D_0__Bacteria;D_1__Firmicutes;D_2__Clostridia;D_3__Clostridiales;D_4__Lachnospiraceae;D_5__uncultured;D_6__Clostridium_sp._Culture-54 | All others | 0.238576 | 0.067338 | 0.584367 | 0.506245 | 0.223595 | 0.477269 | 0.189107 | 0.300991 | 0.372971 | 0.009821 | 0.0016164 | 0.0249358 | 0.249736163 |
| 47 | D_0__Bacteria;D_1__Actinobacteria;D_2__Actinobacteria;D_3__Bifidobacteriales;D_4__Bifidobacteriaceae;D_5__Bifidobacterium;__ | All others | 0.027685 | 0.028751 | 0.013132 | 0.149384 | 0.198588 | 0.004558 | 0.013448 | 0.025556 | 0.004782 | 0.4332781 | 0.6562412 | 1.7324935 | 0.273991234 |
| 48 | D_0__Bacteria;D_1__Firmicutes;D_2__Clostridia;D_3__Clostridiales;D_4__Lachnospiraceae;D_5__GCA-900066575;D_6__uncultured_bacterium | All others | 0.319187 | 0.29432 | 0.429599 | 0.174281 | 0.188291 | 0.375044 | 0.148764 | 0.296732 | 0.245232 | 0.0849225 | 0.1179941 | 0.1474463 | 0.235151095 |
| 49 | D_0__Bacteria;D_1__Firmicutes;D_2__Bacilli;D_3__Bacillales;D_4__Planococcaceae;D_5__Sporosarcina;__ | All others | 0 | 0 | 0 | 0 | 0.000736 | 0 | 2.886199 | 0.018457 | 0 | 0 | 0 | 0 | 0.242115989 |
| 50 | D_0__Bacteria;D_1__Firmicutes;D_2__Clostridia;D_3__Clostridiales;D_4__Ruminococcaceae;D_5__uncultured;D_6__uncultured_Oscillospira_sp. | All others | 0.232062 | 0.350309 | 0.091923 | 0.075522 | 0.275816 | 0.353557 | 0.132795 | 0.169662 | 0.311492 | 0.2380141 | 0.1737584 | 0.1073322 | 0.209353691 |
| 51 | D_0__Bacteria;D_1__Firmicutes;D_2__Clostridia;D_3__Clostridiales;D_4__Lachnospiraceae;D_5__[Eubacterium]_ventriosum_group;__ | All others | 0.247533 | 0.177803 | 0.208234 | 0.340263 | 0.126508 | 0.277376 | 0.1412 | 0.191669 | 0.152331 | 0.201041 | 0.2658908 | 0.1908128 | 0.210055054 |
| 52 | D_0__Bacteria;D_1__Firmicutes;D_2__Clostridia;D_3__Clostridiales;D_4__Ruminococcaceae;D_5__Ruminiclostridium_5;__ | All others | 0.772726 | 0.523572 | 0.166962 | 0.123657 | 0.084584 | 0.213567 | 0.089931 | 0.107193 | 0.101098 | 0.0958989 | 0.0743524 | 0.0542082 | 0.200645713 |
| 53 | D_0__Bacteria;D_1__Bacteroidetes;D_2__Bacteroidia;D_3__Bacteroidales;D_4__Muribaculaceae;D_5__mouse_gut_metagenome;D_6__mouse_gut_metagenome | All others | 0.293945 | 0.391922 | 0.225117 | 0.121167 | 0.033098 | 0.207056 | 0.305934 | 0.43019 | 0.253429 | 0 | 0 | 0.0975747 | 0.196619426 |
| 54 | D_0__Bacteria;D_1__Firmicutes;D_2__Clostridia;D_3__Clostridiales;D_4__Ruminococcaceae;D_5__Ruminococcaceae_UCG-010;D_6__uncultured_organism | All others | 0.228805 | 0.198988 | 0.166962 | 0.206648 | 0.083848 | 0.115899 | 0.061355 | 0.128489 | 0.081289 | 0.3056055 | 0.3774195 | 0.2786301 | 0.186161424 |
| 55 | D_0__Bacteria;D_1__Firmicutes;D_2__Clostridia;D_3__Clostridiales;D_4__Lachnospiraceae;D_5__A2;D_6__uncultured_bacterium | All others | 0.179136 | 0.345013 | 0.091923 | 0.249803 | 0.231686 | 0.050136 | 0.159691 | 0.149786 | 0.358626 | 0.1270949 | 0.1204186 | 0.1095005 | 0.181067724 |
| 56 | D_0__Bacteria;D_1__Firmicutes;D_2__Bacilli;D_3__Bacillales;D_4__Planococcaceae;D_5__Sporosarcina;D_6__uncultured_bacterium | All others | 0 | 0.00227 | 0 | 0 | 0.011033 | 0 | 2.310472 | 0.035494 | 0.004782 | 0.0051993 | 0.0113145 | 0 | 0.198380381 |
| 57 | D_0__Bacteria;D_1__Firmicutes;D_2__Clostridia;D_3__Clostridiales;D_4__Lachnospiraceae;D_5__[Eubacterium]_xylanophilum_group;D_6__uncultured_bacterium | All others | 0.205192 | 0.214876 | 0.377072 | 0.117017 | 0.034569 | 0.014325 | 0.255505 | 0.333646 | 0.390732 | 0.0306183 | 0.1478967 | 0.0216833 | 0.1785943 |
| 58 | D_0__Bacteria;D_1__Firmicutes;D_2__Bacilli;D_3__Bacillales;D_4__Staphylococcaceae;D_5__Jeotgalicoccus;__ | All others | 0 | 0.003783 | 0 | 0 | 0.017652 | 0 | 2.191965 | 0.008519 | 0 | 0.0034662 | 0 | 0.0043367 | 0.185810151 |
| 59 | D_0__Bacteria;D_1__Firmicutes;D_2__Clostridia;D_3__Clostridiales;D_4__Ruminococcaceae;D_5__uncultured;D_6__unidentified | All others | 0.263004 | 0.272379 | 0.745702 | 0.012449 | 0.030156 | 0.032556 | 0.074802 | 0.095834 | 0.27119 | 0.1409598 | 0.0686952 | 0.112753 | 0.176706573 |
| 60 | D_0__Bacteria;D_1__Firmicutes;D_2__Clostridia;D_3__Clostridiales;D_4__Lachnospiraceae;D_5__Lachnospiraceae_NK4A136_group;D_6__uncultured_bacterium | All others | 0.121324 | 0.12938 | 0.184784 | 0.18341 | 0.075758 | 0.04688 | 0.068919 | 0.24775 | 0.124324 | 0.2605445 | 0.3830767 | 0.0325249 | 0.154889525 |
| 61 | D_0__Bacteria;D_1__Firmicutes;D_2__Clostridia;D_3__Clostridiales;D_4__Ruminococcaceae;D_5__Angelakisella;__ | All others | 0.197049 | 0.187639 | 0.222304 | 0.124486 | 0.189762 | 0.213567 | 0.217684 | 0.298861 | 0.170091 | 0.0300406 | 0 | 0.0368616 | 0.157362044 |
| 62 | D_0__Bacteria;D_1__Actinobacteria;D_2__Coriobacteriia;D_3__Coriobacteriales;D_4__Eggerthellaceae;D_5__Enterorhabdus;__ | All others | 0.311045 | 0.217903 | 0.496196 | 0.100419 | 0.045602 | 0.088552 | 0.065557 | 0.078087 | 0.407809 | 0.0635474 | 0.0379844 | 0.0119258 | 0.160385663 |
| 63 | D_0__Bacteria;D_1__Firmicutes;D_2__Clostridia;D_3__Clostridiales;D_4__Ruminococcaceae;D_5__Ruminococcaceae_UCG-004;D_6__uncultured_bacterium | All others | 0.161222 | 0.139972 | 0.222304 | 0.18258 | 0.096352 | 0.212264 | 0.235334 | 0.181021 | 0.148915 | 0.1351828 | 0.0759688 | 0.0715548 | 0.155222514 |
| 64 | D_0__Bacteria;D_1__Firmicutes;D_2__Clostridia;D_3__Clostridiales;D_4__Ruminococcaceae;D_5__Ruminococcaceae_UCG-009;Ambiguous_taxa | All others | 0.184835 | 0.287511 | 0.232621 | 0.078841 | 0.186084 | 0.298212 | 0.08825 | 0.192379 | 0.196049 | 0.0277298 | 0.0266699 | 0.0379457 | 0.153094021 |
| 65 | D_0__Bacteria;D_1__Firmicutes;D_2__Clostridia;D_3__Clostridiales;D_4__Family_XIII;D_5__Family_XIII_AD3011_group;__ | All others | 0.377813 | 0.236818 | 0.138822 | 0.132786 | 0.127979 | 0.074227 | 0.074802 | 0.105773 | 0.157796 | 0.0306183 | 0.1139532 | 0.0314407 | 0.133569051 |
| 66 | D_0__Bacteria;D_1__Firmicutes;D_2__Clostridia;D_3__Clostridiales;D_4__Ruminococcaceae;D_5__Anaerotruncus;D_6__uncultured_bacterium | All others | 0.088754 | 0.118787 | 0.034706 | 0.149384 | 0.249338 | 0.218125 | 0.076483 | 0.091575 | 0.215176 | 0.0699022 | 0.1212268 | 0.0704706 | 0.125327224 |
| 67 | D_0__Bacteria;D_1__Firmicutes;D_2__Clostridia;D_3__Clostridiales;D_4__Family_XIII;D_5__[Eubacterium]_nodatum_group;__ | All others | 0.095268 | 0.052962 | 0.279521 | 0.271381 | 0.163283 | 0.136084 | 0.063036 | 0.076668 | 0.081289 | 0.0895441 | 0.0412171 | 0.125763 | 0.123001218 |
| 68 | D_0__Bacteria;D_1__Firmicutes;D_2__Clostridia;D_3__Clostridiales;D_4__Lachnospiraceae;D_5__Lachnospiraceae_NK4A136_group;D_6__Clostridiales_bacterium_CIEAF_020 | All others | 0.789825 | 0.668841 | 0 | 0 | 0 | 0 | 0 | 0 | 0 | 0.0011554 | 0 | 0 | 0.121651763 |

| | | | | | | | | | | | | | | | |
|---|---|---|---|---|---|---|---|---|---|---|---|---|---|---|---|
| 69 | D_0__Bacteria;D_1__Firmicutes;D_2__Clostridia;D_3__Clostridiales;D_4__Lachnospiraceae;D_5__Marvinbryantia;D_6__uncultured_bacterium | All others | 0.029313 | 0.014376 | 0.027202 | 0.027387 | 0.01471 | 0.026696 | 0.222727 | 0.585655 | 0.012296 | 0.2016187 | 0.1503213 | 0.0357774 | 0.112339808 |
| 70 | D_0__Bacteria;D_1__Firmicutes;D_2__Clostridia;D_3__Clostridiales;D_4__Peptococcaceae;D_5__uncultured;D_6__uncultured_bacterium | All others | 0.145751 | 0.152078 | 0.160396 | 0.114528 | 0.085319 | 0.102877 | 0.08741 | 0.118551 | 0.125007 | 0.0693245 | 0.076777 | 0.125763 | 0.113648376 |
| 71 | D_0__Bacteria;D_1__Firmicutes;D_2__Clostridia;D_3__Clostridiales;D_4__Ruminococcaceae;D_5__Ruminiclostridium_5;Ambiguous_taxa | All others | 0.03257 | 0.04237 | 0.16321 | 0.051454 | 0.036776 | 0.034509 | 0.045386 | 0.092995 | 0.079922 | 0.308494 | 0.2295228 | 0.0585448 | 0.097979508 |
| 72 | D_0__Bacteria;D_1__Firmicutes;D_2__Clostridia;D_3__Clostridiales;D_4__Ruminococcaceae;D_5__Ruminococcaceae_UCG-005;__ | All others | 0.002443 | 0 | 0 | 0.00332 | 0.001471 | 0 | 0.368129 | 0.670841 | 0.003415 | 0.0046216 | 0 | 0.1712979 | 0.10212819 |
| 73 | D_0__Bacteria;D_1__Actinobacteria;D_2__Coriobacteriia;D_3__Coriobacteriales;D_4__Eggerthellaceae;D_5__Enterorhabdus;D_6__mouse_gut_metagenome | All others | 0.119695 | 0.053719 | 0.268265 | 0.170961 | 0.09341 | 0.218125 | 0.066398 | 0.094415 | 0.038937 | 0.0277298 | 0.0202045 | 0.0108416 | 0.09855832 |
| 74 | D_0__Bacteria;D_1__Firmicutes;D_2__Clostridia;D_3__Clostridiales;D_4__Ruminococcaceae;D_5__Ruminococcaceae_UCG-013;D_6__uncultured_rumen_bacterium | All others | 0.078168 | 0.08247 | 0.075039 | 0.151044 | 0.084584 | 0.14585 | 0.062195 | 0.113582 | 0.178288 | 0.0196419 | 0.0145472 | 0.0032525 | 0.084055187 |
| 75 | D_0__Bacteria;D_1__Firmicutes;D_2__Clostridia;D_3__Clostridiales;D_4__Ruminococcaceae;D_5__Ruminococcaceae_NK4A214_group;D_6__uncultured_bacterium | All others | 0.157965 | 0.115004 | 0.155706 | 0.134445 | 0.083848 | 0.118503 | 0.045386 | 0.06176 | 0.068993 | 0.0265744 | 0.0274781 | 0.0422824 | 0.086495503 |
| 76 | D_0__Bacteria;D_1__Firmicutes;D_2__Clostridia;D_3__Clostridiales;D_4__Lachnospiraceae;D_5__Marvinbryantia;Ambiguous_taxa | All others | 0.118881 | 0.060529 | 0.151016 | 0.147724 | 0.076493 | 0.073576 | 0.064717 | 0.097964 | 0.120908 | 0.0340845 | 0.0533398 | 0.006505 | 0.083811461 |
| 77 | D_0__Bacteria;D_1__Actinobacteria;D_2__Coriobacteriia;D_3__Coriobacteriales;D_4__Eggerthellaceae;D_5__Adlercreutzia;D_6__uncultured_bacterium | All others | 0.144123 | 0.094576 | 0.344242 | 0.148554 | 0.02133 | 0.162128 | 0.030257 | 0.040463 | 0.000683 | 0 | 0.0024245 | 0 | 0.082398421 |
| 78 | D_0__Bacteria;D_1__Firmicutes;D_2__Clostridia;D_3__Clostridiales;D_4__Family_XIII;D_5__[Eubacterium]_brachy_group;D_6__uncultured_bacterium | All others | 0.191349 | 0.072634 | 0.232621 | 0.073032 | 0.046337 | 0.044927 | 0.036981 | 0.070988 | 0.093584 | 0.0271521 | 0.031519 | 0.0271041 | 0.079019199 |
| 79 | D_0__Bacteria;D_1__Firmicutes;D_2__Clostridia;D_3__Clostridiales;D_4__Ruminococcaceae;D_5__Ruminococcus_2;__ | All others | 0 | 0 | 0 | 0.160173 | 0 | 0.582099 | 0 | 0.00142 | 0 | 0.0069324 | 0.0622298 | 0 | 0.067737839 |
| 80 | D_0__Bacteria;D_1__Firmicutes;D_2__Erysipelotrichia;D_3__Erysipelotrichales;D_4__Erysipelotrichaceae;D_5__uncultured_bacterium;D_6__uncultured_bacterium | All others | 0.075725 | 0.072634 | 0.126629 | 0.08797 | 0.051486 | 0.040369 | 0.104219 | 0.128489 | 0.073091 | 0.005777 | 0.0266699 | 0.0021683 | 0.06626907 |
| 81 | D_0__Bacteria;D_1__Firmicutes;D_2__Clostridia;D_3__Clostridiales;D_4__Ruminococcaceae;D_5__Ruminococcaceae_UCG-014;D_6__uncultured_bacterium | All others | 0.16855 | 0.255733 | 0.114435 | 0 | 0.001471 | 0.003256 | 0.002521 | 0 | 0.002049 | 0.0271521 | 0.0880915 | 0.0271041 | 0.057530268 |
| 82 | D_0__Bacteria;D_1__Actinobacteria;D_2__Coriobacteriia;D_3__Coriobacteriales;D_4__Eggerthellaceae;D_5__Enterorhabdus;D_6__uncultured_bacterium | All others | 0.11481 | 0.125597 | 0.066597 | 0.024897 | 0.019859 | 0.016929 | 0.010926 | 0.017037 | 0.017761 | 0.0947435 | 0.1147614 | 0.0162625 | 0.05334834 |
| 83 | D_0__Bacteria;D_1__Firmicutes;D_2__Clostridia;D_3__Clostridiales;D_4__Lachnospiraceae;D_5__A2;__ | All others | 0.086311 | 0.0401 | 0.484003 | 0.005809 | 0.005149 | 0 | 0 | 0.008519 | 0.003415 | 0.0306183 | 0.0282863 | 0.0401141 | 0.061027018 |
| 84 | D_0__Bacteria;D_1__Proteobacteria;D_2__Gammaproteobacteria;D_3__Betaproteobacteriales;D_4__Burkholderiaceae;D_5__Parasutterella;__ | All others | 0.140866 | 0.074147 | 0.213862 | 0.028217 | 0.02133 | 0.037765 | 0.028576 | 0.070988 | 0.044401 | 0 | 0 | 0 | 0.055012701 |
| 85 | D_0__Bacteria;D_1__Actinobacteria;D_2__Coriobacteriia;D_3__Coriobacteriales;D_4__Eggerthellaceae;__;__ | All others | 0.020356 | 0.035561 | 0 | 0.066393 | 0.035305 | 0.062507 | 0.130274 | 0.108612 | 0.056014 | 0.0179088 | 0.0121227 | 0.0216833 | 0.047228046 |
| 86 | D_0__Bacteria;D_1__Firmicutes;D_2__Clostridia;D_3__Clostridiales;D_4__Lachnospiraceae;D_5__Lachnospiraceae_FCS020_group;D_6__uncultured_bacterium | All others | 0.043155 | 0.046153 | 0.030016 | 0.00332 | 0.019123 | 0.031254 | 0.031938 | 0.054661 | 0.090169 | 0.0716353 | 0.0210126 | 0.0303566 | 0.039399442 |
| 87 | D_0__Bacteria;D_1__Firmicutes;D_2__Clostridia;D_3__Clostridiales;D_4__Clostridiales_vadinBB60_group;Ambiguous_taxa;Ambiguous_taxa | All others | 0.030127 | 0.200501 | 0.011256 | 0 | 0.009562 | 0 | 0.005883 | 0.009938 | 0.064894 | 0.0485271 | 0.0646543 | 0.0401141 | 0.040454768 |
| 88 | D_0__Bacteria;D_1__Firmicutes;D_2__Clostridia;D_3__Clostridiales;D_4__Ruminococcaceae;D_5__Oscillibacter;__ | All others | 0.024428 | 0.062798 | 0.267327 | 0.004979 | 0.05737 | 0.021487 | 0.004202 | 0.00071 | 0.008197 | 0.0103987 | 0.022629 | 0.0151783 | 0.041642046 |
| 89 | D_0__Bacteria;D_1__Proteobacteria;D_2__Gammaproteobacteria;D_3__Pseudomonadales;D_4__Pseudomonadaceae;D_5__Pseudomonas;__ | All others | 0.013028 | 0.011349 | 0.019698 | 0.019918 | 0.025007 | 0.00586 | 0.005043 | 0.009938 | 0.017761 | 0.0387062 | 0.2707399 | 0.0195149 | 0.038046914 |
| 90 | D_0__Bacteria;D_1__Firmicutes;D_2__Clostridia;D_3__Clostridiales;D_4__Ruminococcaceae;D_5__GCA-900066225;Ambiguous_taxa | All others | 0.072468 | 0.057502 | 0.045961 | 0.072202 | 0.047808 | 0.067065 | 0 | 0 | 0.064894 | 0.0115541 | 0 | 0 | 0.036621329 |
| 91 | D_0__Bacteria;D_1__Firmicutes;D_2__Clostridia;D_3__Clostridiales;D_4__Ruminococcaceae;D_5__uncultured;D_6__uncultured_bacterium | All others | 0.017099 | 0.012862 | 0.03283 | 0.015768 | 0.027949 | 0.117201 | 0.068919 | 0.075248 | 0.041669 | 0 | 0.0105063 | 0.006505 | 0.035546444 |
| 92 | D_0__Bacteria;D_1__Firmicutes;D_2__Clostridia;D_3__Clostridiales;D_4__Ruminococcaceae;D_5__GCA-900066225;D_6__uncultured_bacterium | All others | 0.058626 | 0.08247 | 0.094737 | 0.012449 | 0.010297 | 0.007162 | 0.024374 | 0.031945 | 0.050549 | 0.0265744 | 0.0290944 | 0.0184308 | 0.037225738 |
| 93 | D_0__Bacteria;D_1__Firmicutes;D_2__Clostridia;D_3__Clostridiales;D_4__Lachnospiraceae;D_5__Roseburia;D_6__uncultured_bacterium | All others | 0.052112 | 0.024211 | 0.00469 | 0.015768 | 0.02133 | 0.064461 | 0.028576 | 0.044723 | 0.021176 | 0.0121318 | 0.1074878 | 0.0314407 | 0.035675633 |
| 94 | D_0__Bacteria;D_1__Firmicutes;D_2__Clostridia;D_3__Clostridiales;D_4__Defluviitaleaceae;D_5__Defluviitaleaceae_UCG-011;__ | All others | 0 | 0 | 0 | 0.032366 | 0.07208 | 0 | 0.030257 | 0.057501 | 0.045084 | 0.0647029 | 0.054148 | 0.0292724 | 0.032117668 |
| 95 | D_0__Bacteria;D_1__Firmicutes;D_2__Bacilli;D_3__Lactobacillales;D_4__Streptococcaceae;D_5__Streptococcus;__ | All others | 0 | 0 | 0.002814 | 0.043155 | 0.062518 | 0 | 0.027736 | 0.070279 | 0.049183 | 0.041017 | 0 | 0.0227674 | 0.026622452 |
| 96 | D_0__Bacteria;D_1__Firmicutes;D_2__Clostridia;D_3__Clostridiales;D_4__Lachnospiraceae;D_5__Acetatifactor;Ambiguous_taxa | All others | 0.088754 | 0.081714 | 0.033768 | 0.016598 | 0.008091 | 0.024742 | 0.011767 | 0.022716 | 0.028007 | 0.0046216 | 0.0024245 | 0.0108416 | 0.027836984 |
| 97 | D_0__Bacteria;D_1__Firmicutes;D_2__Clostridia;D_3__Clostridiales;D_4__Lachnospiraceae;D_5__Lachnospiraceae_NK4A136_group;Ambiguous_taxa | All others | 0.07084 | 0.071878 | 0.034706 | 0.009129 | 0.011768 | 0 | 0.026055 | 0.022716 | 0.079239 | 0 | 0 | 0.0043367 | 0.027555625 |
| 98 | D_0__Bacteria;D_1__Firmicutes;D_2__Clostridia;D_3__Clostridiales;D_4__Lachnospiraceae;D_5__GCA-900066575;__ | All others | 0.0114 | 0.006809 | 0 | 0.032366 | 0.022065 | 0.013022 | 0.033619 | 0.051822 | 0.012979 | 0.0358177 | 0.0193963 | 0.0748073 | 0.026175326 |
| 99 | D_0__Bacteria;D_1__Firmicutes;D_2__Clostridia;D_3__Clostridiales;D_4__Clostridiales_vadinBB60_group;__;__ | All others | 0.008957 | 0.043127 | 0.008442 | 0 | 0.019859 | 0.005209 | 0 | 0.004259 | 0.007514 | 0.0346622 | 0.0274781 | 0.0563765 | 0.017990269 |
| 100 | D_0__Bacteria;D_1__Firmicutes;D_2__Clostridia;D_3__Clostridiales;D_4__Ruminococcaceae;D_5__Ruminococcaceae_UCG-013;__ | All others | 0.024428 | 0.010592 | 0.031892 | 0.008299 | 0.005149 | 0.012371 | 0.010926 | 0.023426 | 0 | 0.0127095 | 0.0598052 | 0.0162625 | 0.01798835 |
| 101 | D_0__Bacteria;D_1__Firmicutes;D_2__Bacilli;D_3__Lactobacillales;D_4__Enterococcaceae;D_5__Enterococcus;__ | All others | 0.034199 | 0.018915 | 0.005628 | 0.077182 | 0.028685 | 0.020836 | 0 | 0.00213 | 0.005465 | 0.0069324 | 0.0072736 | 0 | 0.017270377 |
| 102 | D_0__Bacteria;D_1__Proteobacteria;D_2__Alphaproteobacteria;D_3__Rhizobiales;D_4__Beijerinckiaceae;D_5__Methylobacterium;__ | All others | 0.002443 | 0.003026 | 0 | 0.006639 | 0 | 0 | 0 | 0 | 0 | 0.0190642 | 0.153554 | 0 | 0.015393889 |
| 103 | D_0__Bacteria;D_1__Firmicutes;D_2__Clostridia;D_3__Clostridiales;D_4__Ruminococcaceae;D_5__Harryflintia;D_6__uncultured_bacterium | All others | 0.006514 | 0.024968 | 0.01407 | 0.007469 | 0.016917 | 0.031254 | 0.010086 | 0.018457 | 0.022542 | 0.009821 | 0 | 0.0075891 | 0.014140545 |
| 104 | D_0__Bacteria;D_1__Firmicutes;D_2__Clostridia;D_3__Clostridiales;D_4__Ruminococcaceae;D_5__Ruminococcaceae_UCG-005;Ambiguous_taxa | All others | 0.008143 | 0.015889 | 0.016884 | 0.009129 | 0.005884 | 0.001953 | 0.005883 | 0.015617 | 0.014345 | 0.0398616 | 0.013739 | 0.0173466 | 0.013722885 |
| 105 | D_0__Bacteria;D_1__Firmicutes;D_2__Clostridia;D_3__Clostridiales;D_4__Ruminococcaceae;D_5__Anaerotruncus;D_6__uncultured_organism | All others | 0 | 0 | 0 | 0.050625 | 0.077964 | 0.035811 | 0 | 0 | 0 | 0 | 0 | 0 | 0.013700009 |
| 106 | D_0__Bacteria;D_1__Firmicutes;D_2__Clostridia;D_3__Clostridiales;D_4__Lachnospiraceae;D_5__Roseburia;__ | All others | 0.0114 | 0.010592 | 0.043148 | 0.024897 | 0.015446 | 0.013673 | 0.005043 | 0.022716 | 0.011613 | 0.0028885 | 0.0080818 | 0.0032525 | 0.014395889 |
| 107 | D_0__Bacteria;D_1__Firmicutes;D_2__Clostridia;D_3__Clostridiales;D_4__Clostridiaceae_1;D_5__Candidatus_Arthromitus;D_6__uncultured_bacterium | All others | 0.122952 | 0.037074 | 0 | 0 | 0 | 0 | 0 | 0 | 0 | 0 | 0 | 0 | 0.013335492 |
| 108 | D_0__Bacteria;D_1__Firmicutes;D_2__Clostridia;D_3__Clostridiales;D_4__Ruminococcaceae;D_5__Butyricicoccus;D_6__uncultured_bacterium | All others | 0.004071 | 0.001513 | 0 | 0.008299 | 0 | 0.033207 | 0.014288 | 0.014908 | 0.021859 | 0.0184865 | 0.0169718 | 0.006505 | 0.011675722 |
| 109 | D_0__Bacteria;D_1__Tenericutes;D_2__Mollicutes;D_3__Mollicutes_RF39;__;__;__ | All others | 0.030942 | 0.033291 | 0.021574 | 0 | 0 | 0 | 0 | 0 | 0 | 0.0127095 | 0.0193963 | 0.0119258 | 0.010819804 |
| 110 | D_0__Bacteria;D_1__Firmicutes;D_2__Clostridia;D_3__Clostridiales;D_4__Family_XIII;D_5__Family_XIII_AD3011_group;D_6__uncultured_bacterium | All others | 0.013842 | 0.027238 | 0.007504 | 0.009129 | 0.011768 | 0.010418 | 0.009245 | 0.009938 | 0.007514 | 0.0034662 | 0 | 0.0021683 | 0.009352616 |
| 111 | D_0__Bacteria;D_1__Firmicutes;D_2__Clostridia;D_3__Clostridiales;D_4__Lachnospiraceae;D_5__ASF356;D_6__uncultured_bacterium | All others | 0 | 0.012106 | 0 | 0 | 0 | 0 | 0.010926 | 0.024846 | 0.002049 | 0 | 0.054148 | 0.0119258 | 0.009666745 |
| 112 | D_0__Bacteria;D_1__Firmicutes;D_2__Clostridia;D_3__Clostridiales;D_4__Clostridiales_vadinBB60_group;D_5__uncultured_Clostridia_bacterium;D_6__uncultured_Clostridia_bact | All others | 0 | 0.015889 | 0.001876 | 0 | 0 | 0 | 0.004202 | 0 | 0 | 0.0225305 | 0.0266699 | 0.0477032 | 0.009905889 |
| 113 | D_0__Bacteria;D_1__Firmicutes;D_2__Clostridia;D_3__Clostridiales;D_4__Ruminococcaceae;D_5__Ruminococcus_1;__ | All others | 0.004071 | 0.006809 | 0.005628 | 0 | 0 | 0.002604 | 0 | 0.005679 | 0.004782 | 0.0190642 | 0.0468744 | 0.0032525 | 0.008230416 |
| 114 | D_0__Bacteria;D_1__Firmicutes;D_2__Bacilli;D_3__Lactobacillales;D_4__Lactobacillaceae;D_5__Lactobacillus;D_6__Lactobacillus_reuteri | All others | 0 | 0 | 0 | 0.08963 | 0.018388 | 0 | 0 | 0 | 0 | 0 | 0 | 0 | 0.009001503 |
| 115 | D_0__Bacteria;D_1__Tenericutes;D_2__Mollicutes;D_3__Mollicutes_RF39;D_4__uncultured_bacterium;D_5__uncultured_bacterium;D_6__uncultured_bacterium | All others | 0.007328 | 0.027994 | 0.013132 | 0.013279 | 0.022065 | 0 | 0 | 0 | 0 | 0 | 0 | 0 | 0.006983205 |
| 116 | D_0__Bacteria;D_1__Firmicutes;D_2__Clostridia;D_3__Clostridiales;D_4__Lachnospiraceae;D_5__uncultured;D_6__Clostridium_sp._Culture-27 | All others | 0 | 0 | 0.04221 | 0.009129 | 0.005149 | 0.003256 | 0 | 0.00284 | 0 | 0.005777 | 0.0121227 | 0.0054208 | 0.007158565 |
| 117 | D_0__Bacteria;D_1__Proteobacteria;D_2__Gammaproteobacteria;D_3__Enterobacteriales;D_4__Enterobacteriaceae;D_5__Escherichia-Shigella;__ | All others | 0.001629 | 0.003783 | 0.003752 | 0.00249 | 0 | 0 | 0.001681 | 0 | 0 | 0.005777 | 0.0573807 | 0 | 0.006374326 |
| 118 | D_0__Bacteria;D_1__Firmicutes;D_2__Clostridia;D_3__Clostridiales;D_4__Lachnospiraceae;D_5__Acetatifactor;__ | All others | 0.007328 | 0.010592 | 0.045961 | 0.00249 | 0.005884 | 0.001953 | 0.001681 | 0.00284 | 0 | 0 | 0 | 0 | 0.006560827 |
| 119 | D_0__Bacteria;D_1__Firmicutes;D_2__Clostridia;D_3__Clostridiales;D_4__Lachnospiraceae;D_5__uncultured;D_6__uncultured_Clostridiales_bacterium | All others | 0.008143 | 0.014376 | 0.027202 | 0 | 0 | 0 | 0 | 0.006389 | 0 | 0 | 0.0096981 | 0 | 0.005483905 |
| 120 | D_0__Bacteria;D_1__Firmicutes;D_2__Clostridia;D_3__Clostridiales;D_4__Lachnospiraceae;D_5__Lachnospiraceae_UCG-001;D_6__uncultured_bacterium | All others | 0.001629 | 0.00454 | 0.003752 | 0.00332 | 0.002207 | 0 | 0.001681 | 0.011358 | 0.005465 | 0.0092433 | 0.0048491 | 0 | 0.004003542 |
| 121 | D_0__Bacteria;D_1__Firmicutes;D_2__Clostridia;D_3__Clostridiales;D_4__Lachnospiraceae;D_5__Lachnospiraceae_NK4A136_group;D_6__Lachnospiraceae_bacterium_A4 | All others | 0 | 0 | 0 | 0 | 0 | 0 | 0 | 0.004259 | 0 | 0 | 0.0064654 | 0.0140941 | 0.002068239 |
| 122 | D_0__Bacteria;D_1__Firmicutes;D_2__Clostridia;D_3__Clostridiales;D_4__Lachnospiraceae;D_5__Lachnospira;__ | All others | 0 | 0 | 0 | 0 | 0 | 0 | 0.005883 | 0.009228 | 0 | 0 | 0 | 0.0021683 | 0.001440014 |
| 123 | D_0__Bacteria;D_1__Firmicutes;D_2__Clostridia;D_3__Clostridiales;D_4__Ruminococcaceae;D_5__Ruminiclostridium_5;D_6__uncultured_organism | All others | 0.003257 | 0.006053 | 0 | 0 | 0 | 0 | 0 | 0 | 0 | 0.0023108 | 0 | 0 | 0.00096839 |
| 124 | D_0__Bacteria;D_1__Firmicutes;D_2__Erysipelotrichia;D_3__Erysipelotrichales;D_4__Erysipelotrichaceae;D_5__uncultured;D_6__uncultured_bacterium | All others | 0 | 0 | 0 | 0 | 0 | 0 | 0 | 0 | 0 | 0 | 0.0105063 | 0.0021683 | 0.001056221 |
| 125 | D_0__Bacteria;D_1__Firmicutes;D_2__Clostridia;D_3__Clostridiales;D_4__Ruminococcaceae;D_5__Ruminococcaceae_UCG-014;D_6__uncultured_rumen_bacterium | All others | 0 | 0 | 0 | 0 | 0 | 0 | 0 | 0.007099 | 0.001366 | 0 | 0 | 0 | 0.00070542 |
| 126 | D_0__Bacteria;D_1__Firmicutes;D_2__Clostridia;D_3__Clostridiales;D_4__Ruminococcaceae;D_5__Ruminiclostridium_6;D_6__uncultured_bacterium | All others | 0 | 0 | 0 | 0 | 0 | 0 | 0 | 0 | 0 | 0 | 0.0072736 | 0 | 0.000606134 |
| 127 | D_0__Bacteria;D_1__Firmicutes;D_2__Clostridia;D_3__Clostridiales;D_4__Lachnospiraceae;D_5__Tyzzerella_3;D_6__uncultured_bacterium | All others | 0 | 0 | 0 | 0 | 0 | 0.003907 | 0 | 0 | 0 | 0 | 0 | 0 | 0.000325559 |
| 128 | D_0__Bacteria;D_1__Firmicutes;D_2__Clostridia;D_3__Clostridiales;D_4__Ruminococcaceae;D_5__Ruminiclostridium_6;__ | All others | 0 | 0 | 0 | 0 | 0.001471 | 0 | 0 | 0 | 0 | 0 | 0 | 0 | 0.000122585 |
| 129 | D_0__Bacteria;D_1__Tenericutes;D_2__Mollicutes;D_3__Anaeroplasmatales;D_4__Anaeroplasmataceae;D_5__Anaeroplasma;D_6__uncultured_bacterium | All others | 0 | 0 | 0 | 0 | 0 | 0 | 0 | 0.00142 | 0 | 0 | 0 | 0 | 0.000118314 |
| 130 | D_0__Bacteria;D_1__Firmicutes;D_2__Erysipelotrichia;D_3__Erysipelotrichales;D_4__Erysipelotrichaceae;__;__ | All others | 0 | 0 | 0 | 0 | 0 | 0 | 0 | 0 | 0 | 0 | 0 | 0 | 0 |
| | | **TOTAL (each sample)** | 100 | 100 | 100 | 100 | 100 | 100 | 100 | 100 | 100 | 100 | 100 | 100 | |

Supplementary Table 2: Proximal nutritional analysis of the modified chow diets consumed in the study.

| Analyte / Diet | Frankfurter | Pork | Sausage | Control |
|---|---|---|---|---|
| Sodium nitrate mg/Kg | 24.6 | 31.9 | <21 | 25.3 |
| Potassium nitrate mg/Kg | 29.3 | 38 | <25 | 30.1 |
| Sodium nitrite mg/Kg | <22.5 | <22.5 | <22.5 | <22.5 |
| Potassium nitrite mg/Kg | <27.7 | <27.7 | <27.7 | <27.7 |
| Fat g/100g | 8.4 | 6.2 | 8.2 | 3.5 |
| Moisture g/100g | 9.3 | 8.3 | 8.3 | 7 |
| Ash g/100g | 2.9 | 2.7 | 2.7 | 2.6 |
| Sodium g/100g | 0.26 | 0.11 | 0.21 | 0.11 |
| Sodium chloride g/100g | 0.65 | 0.28 | 0.53 | 0.28 |
| Total sugars g/100g | 44.6 | 47.4 | 46 | 51.7 |
| Nitrogen (mg/Kg) g/100g | 3 | 3.41 | 2.98 | 2.85 |
| Protein g/100g | 18.8 | 21.3 | 18.6 | 17.8 |
| Monounsaturated fatty acids g/100g | 2.35 | 3.13 | 3.84 | 1.81 |
| Polyunsaturated fatty acids g/100g | 2.5 | 0.25 | 0.27 | 0.25 |
| Saturated fatty acids g/100g | 2.18 | 2.55 | 3.73 | 1.28 |
| Dietary fibre g/100g | 5 | 6.3 | 6.3 | 7.1 |
| Carbohydrate g/100g | 55.6 | 55.2 | 55.9 | 62 |
| Energy kcal/100g | 383 | 374 | 385 | 362 |
